# Genome-wide transcriptomics identifies an early preclinical signature of prion infection

**DOI:** 10.1101/2020.01.10.901637

**Authors:** Silvia Sorce, Mario Nuvolone, Giancarlo Russo, Andra Chincisan, Daniel Heinzer, Merve Avar, Manuela Pfammatter, Petra Schwarz, Mirzet Delic, Simone Hornemann, Despina Sanoudou, Claudia Scheckel, Adriano Aguzzi

## Abstract

The clinical course of prion diseases is accurately predictable despite long latency periods, suggesting that prion pathogenesis is driven by precisely timed molecular events. We constructed a searchable genome-wide atlas of mRNA abundance, splicing and editing alterations during the course of disease in prion-inoculated mice. Prion infection induced transient changes in mRNA abundance and processing already at eight weeks post inoculation, well ahead of any neuropathological and clinical signs. In contrast, microglia-enriched genes displayed an increase simultaneous with the appearance of clinical symptoms, whereas neuronal-enriched transcripts remained unchanged until the very terminal stage of disease. This suggests that glial pathophysiology, rather than neuronal demise, represents the final driver of disease. The administration of young plasma attenuated the occurrence of early mRNA abundance alterations and delayed symptoms in the terminal phase of the disease. The early onset of prion-induced molecular changes might thus point to novel biomarkers and potential interventional targets.

## Main

After the onset of clinical signs, prion diseases typically progress very rapidly to a terminal stage, which is characterized by micro- and astrogliosis, vacuolation and neuronal loss. Sporadic Creutzfeldt-Jakob disease (CJD), the most common human prion disease, can lead to death within months of symptom onset^1^. Conversely, prion infections are often characterized by very long incubation times: iatrogenic cases of CJD after administration of prion-contaminated growth hormone display incubation times of >20 years^2^ and Kuru, an acquired form of prion disease, was reported to arise decades after consumption of contaminated materials^3^. The extraordinary duration of the latency phase has led to the presumption that CJD may be caused by a “slow virus”. Although this hypothesis was dismissed^4^, prion pathogenesis is initially insidious and only acquires a rapid rate of progression in the late stages of disease.

The seemingly incongruous combination of a very long latency and a rapidly progressing clinical disease can be reproduced in mouse models of prion infection^5^. After inoculation of prion-containing brain homogenate, wild-type mice experience an incubation period of several months followed by rapidly progressive neurological dysfunction. It was suggested that prion replication occurs without causing any neuronal damage until a plateau level of infectious particles is reached, whereas neurotoxicity arises because of a toxic form of PrP named “PrP^L^“^6–8^. However, no physical evidence for the existence of “PrP^L^“ has ever come forth, and these experimental observations can be explained by alternative models. For example, small numbers of prion seeds may cause early molecular alterations that elicit late-onset clinical signs. This question may be addressed by creating longitudinal maps of the transcriptional equivalents of neurotoxicity from the administration of prions to the development of terminal disease.

Transcriptional maps of prion-inoculated mice were previously performed using microarrays^9–12^, and more recently also by RNA sequencing (RNAseq)^13^. Most of these studies analyzed whole-brain homogenates and have focused on alterations occurring in the late phase of the disease. Yet no antiprion compounds could reverse the progression of disease^14^, perhaps because intervention was too late. Instead, focusing on the preclinical changes occurring in the affected brain may uncover alternative targets for early diagnosis and therapeutic intervention in pre-symptomatic subjects. Here we performed a comprehensive analysis of transcriptional alterations in the hippocampus of prion-infected mice over time. We identified an unexpected wealth of changes in the early phase of prion replication, long before any clinical signs of disease. Neuronal expression changes became evident only at the terminal stage of the disease, whereas the appearance of clinical symptoms coincided with microglia alterations. Prion-induced molecular changes were largely unaffected by ageing, yet the administration of young plasma attenuated early prion-induced changes and improved the health span of diseased mice.

## Results

### Identification of mRNA expression changes during prion disease

To identify molecular alterations associated with the progression of prion disease, we administered intracerebrally RML6 prions or non-infectious brain homogenate (NBH, for control) to C57BL/6 mice (Fig. 1a). Mice were sacrificed at 4, 8, 12, 14, 16, 18 and 20 wpi (weeks post inoculation), as well as the terminal stage (the last time point during disease progression when mice can be humanely euthanized). We specifically focused on the hippocampal region, which plays a central role in memory formation and consolidation and is strongly affected in multiple neurodegenerative disorders including prion disease. RNA from ipsi- and contralateral hippocampi was extracted and subjected to RNA sequencing (*n*=3 per prion and control samples at each time point except for two control samples at 20 wpi; Supplementary Files 1 and 2). All gene expression profiles during disease progression can be visualized and downloaded at PrionRNAseqDatabase (login credentials: username: PrionAG; password: rnarna).

**Fig. 1:**
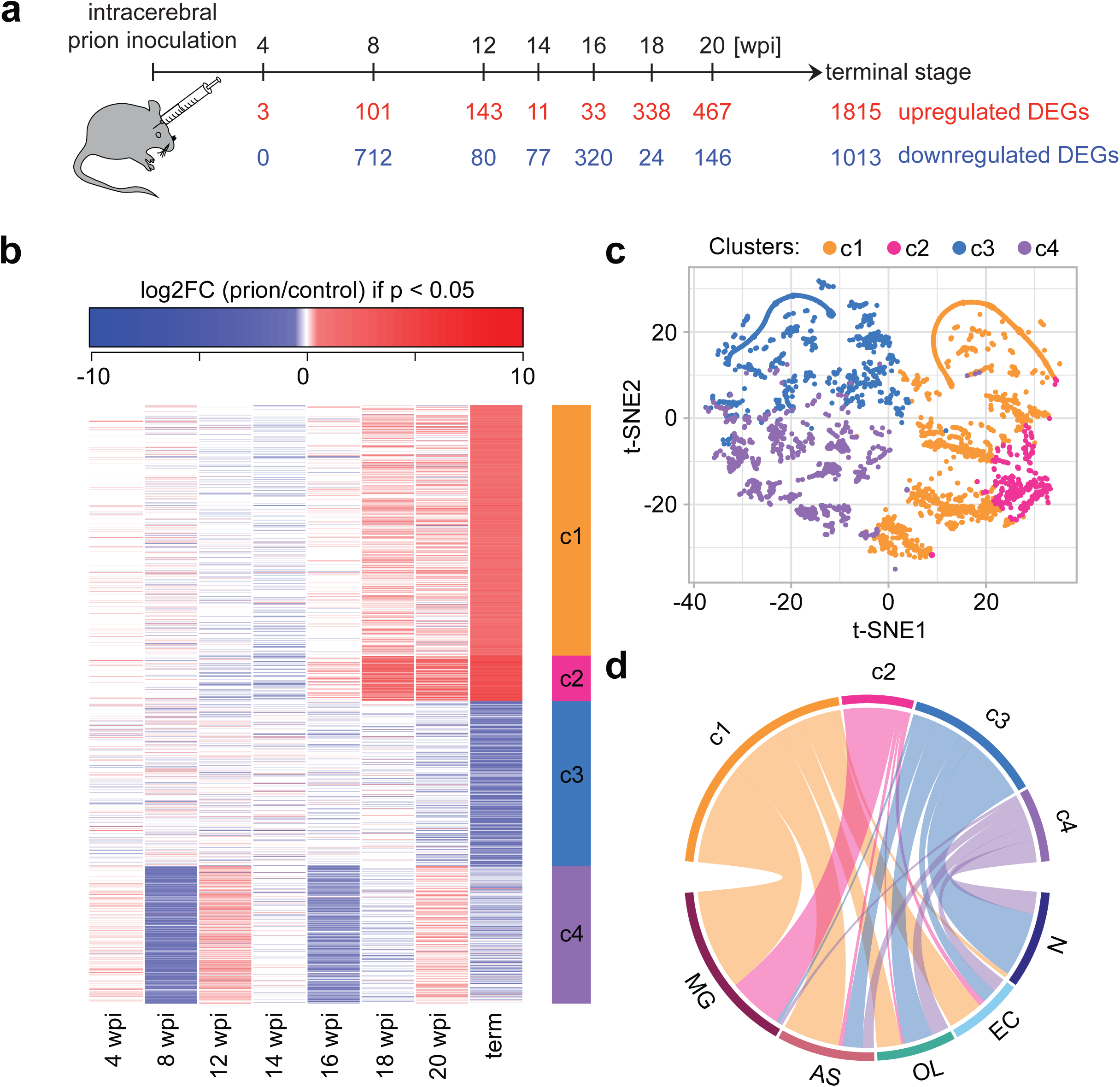
Identification of differentially expressed genes (DEGs) during prion disease progression. **a**, Timeline of prion inoculations. Numbers of upregulated and downregulated DEGs (|log_2_FC| > 0.5 and FDR < 0.05) are indicated. Several DEGs were up- and downregulated at different time points. **b**, Heatmap displaying the log_2_FC of 3,723 genes that are differentially expressed (|log_2_FC| > 0.5 and FDR < 0.05) at least at one time point. Only log_2_FC values with p < 0.05 are colored. Unsupervised kmeans clustering (k = 4 clusters) identified four coherent patterns (c1-c4) of log_2_FC oscillations over time (right side bar) **c**, t-distributed stochastic neighbor embedding (t-SNE) plot (n components = 2, perplexity = 50) visualizing the separation of DEGs into four clusters. **d**, Circos plot summarizing the cell type-enriched genes within each cluster. Clusters are identified by the same colors in panels b-d.

Hierarchical clustering analysis of total transcriptomes revealed a progressive segregation of prion-inoculated and control mice starting at 18 wpi (Supplementary Fig. 1a). However, principal component analysis (PCA) showed a separation of the two groups already at 8 wpi (Supplementary Fig. 1b). Next, we identified 3,723 differentially expressed genes (DEGs) between prion-infected and control mice (absolute log_2_ fold changes |log_2_FC| > 0.5 and false discovery rate (FDR) < 0.05 at least at one of the time points; Fig. 1a-b; Supplementary File 3). Consistent with the hierarchical clustering and the PCA, 813 genes changed at 8 wpi and the number of DEGs gradually increased during the later timepoints (Fig. 1a).

Unsupervised kmeans clustering (k = 4 clusters; log_2_FC with p ≥ 0.05 were set to 0) identified mRNAs with distinct expression patterns during disease progression (Fig. 1b; only log_2_FC with p < 0.05 are colored). The clusters were further visualized with a t-distributed stochastic neighbor embedding (t-SNE) of the log_2_FC (Fig. 1c; log_2_FC with p ≥ 0.05 were set to 0). Clusters 1 and 3 correspond to genes that increase or decrease at the terminal stage only. In contrast, cluster 2 genes started to increase already at 16-18 weeks, and genes in cluster 4 showed an oscillating pattern, decreasing at 8 wpi, 16 wpi and the terminal stage, but recovering in between. Intersecting our data with an atlas of the most enriched transcripts in neurons (N), microglia (MG), astrocytes (AS), oligodendrocytes (OL) and vascular cells (VC)^15^, revealed that the different clusters are populated by distinct cell types. While clusters 2 and 3 consist predominantly of microglia and neuronal genes, respectively, clusters 1 and 4 contain genes enriched in multiple cell types (Fig. 1d). This observation is further substantiated by heatmaps of genes distinctive to just one cell type (Supplementary Fig. 2a). Neuronal genes almost exclusively belonged to clusters 3 and 4, and corresponded to decreasing genes, whereas microglia genes were essentially all increasing and were contained in clusters 1 and 2. At the terminal stage, these patterns may partly reflect shifts in the cellular composition of the brain, with both microglia activation and neuronal loss being hallmarks of prion disease.

### mRNA expression changes become evident in the early disease phase

The most conspicuous gene expression changes arose at the terminal time point (|log_2_FC| > 0.5 and p < 0.05), yet almost half of DEGs were shared between two or more time points (Supplementary Fig. 2b). Multiple patterns of gene expression changes became already evident in the cluster analysis (Fig. 1b). To identify monotonic gene expression patterns, we selected terminal DEGs showing a consistent trend (|log_2_FC| > 0.5 and p value < 0.05) at consecutive time points. Of 1,872 terminally upregulated DEGs, 1, 87, 440 and 632 genes started to be monotonically upregulated at 14, 16, 18, and 20wpi, respectively. Conversely, of 1,081 downregulated DEGs, only 0, 1, 6, and 64 genes showed a continuous decrease at those time points (Fig. 2a). This illustrates that a monotonic pattern is specific to upregulated genes and becomes evident at 16wpi. We further analyzed if these monotonic DEGs (mDEGs) were enriched for certain cell types. Consistent with the cluster analysis, upregulated mDEGs consisted mostly of microglia-enriched and to a lesser extent astrocyte-enriched transcripts (Supplementary Fig. 2c). Moreover, we found that the top enriched GO terms in terminally upregulated genes were shared between mDEGs at 20, 18 and 16 wpi (Fig. 2b; Supplementary File 4). GO terms related to vacuolation (a defining hallmark of prion diseases) started to be enriched at 18 wpi (Supplementary File 5). Both microgliosis and vacuolation are characteristics of prion disease. Our data illustrates that both characteristics also become evident on the RNA expression level, months prior to the terminal stage of the disease. A further feature of human prion diseases is the profound loss of neurons. Yet only a minority (n = 64) of the 1081 terminally downregulated DEGs decreased already at 20 wpi (Supplementary Fig. 2d), and GO terms related to ion channels, synaptic transmission and neuron projection were only enriched among terminally downregulated DEGs (Supplementary File 6).

**Fig. 2:**
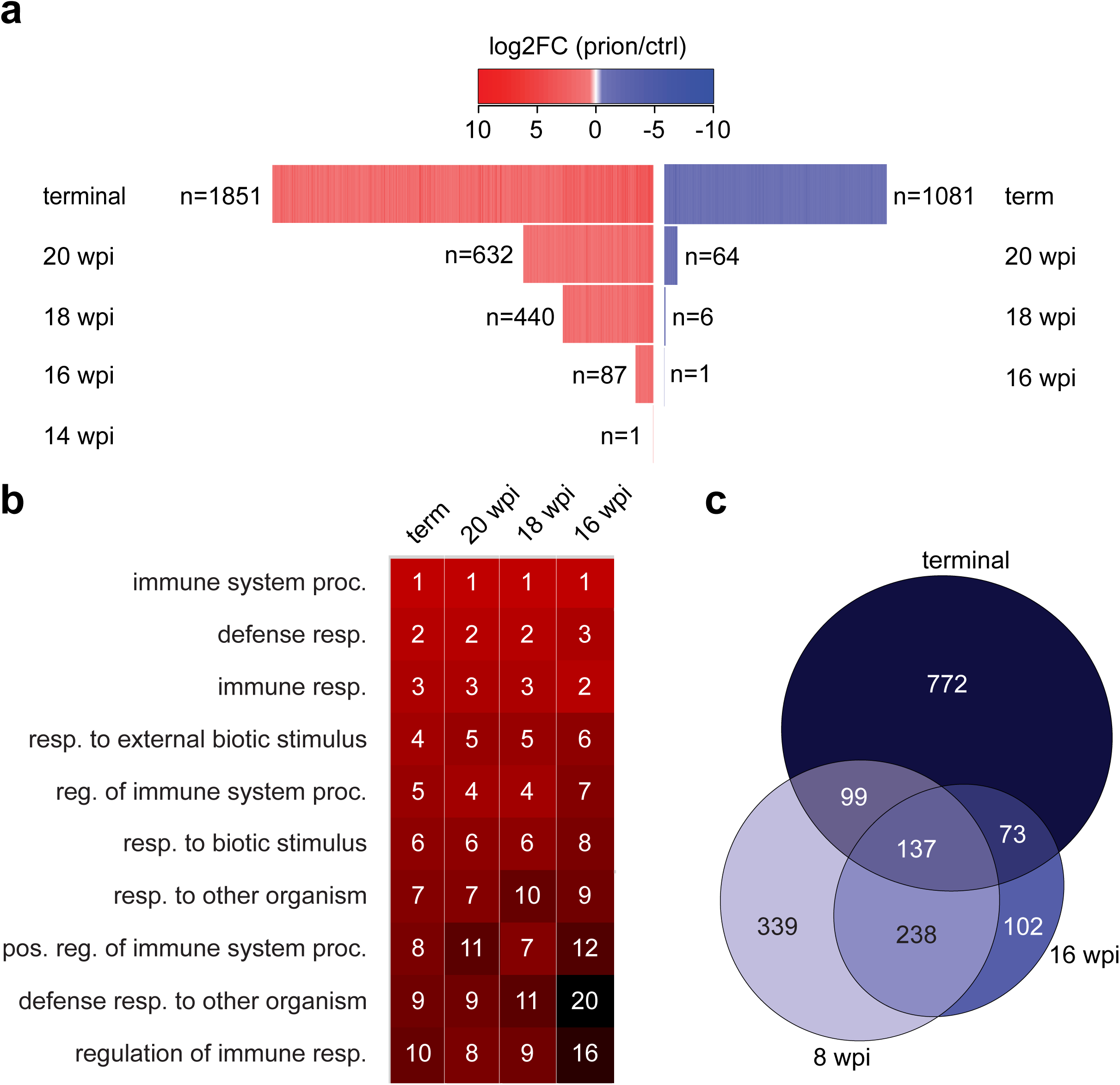
Consistent mRNA expression changes become evident in the early disease phase. **a-b**, **a**, Heatmap displaying the log_2_FC of monotonic DEGs (mDEGs), defined as genes changing with a similar trend (|log_2_FC| > 0.5 and p < 0.05) at consecutive time points until terminal disease. n: number of mDEGs at each time point. **b,** Heatmap displaying the top 10 GO terms enriched among terminally upregulated genes and their enrichment in mDEGs at 20, 18 and 16 wpi. Number and colors represent the ranking (high = red, black = low) of the respective GO terms. **c**, Overlap of downregulated genes at 8 wpi, 16 wpi and at the terminal stage.

We further focused on the oscillating DEGs associated with cluster 4 genes and assessed the overlap between genes downregulated at 8 wpi, 16 wpi and the terminal stage. 813 DEGs showed a decrease (with p < 0.05) already at 8 wpi although neither clinical nor histological changes are detectable at this time point (see below). The 813 genes showed a minor enrichment of neuronal genes (Supplementary Fig. 2e) and were characterized by GO terms related to extracellular matrix, cell adhesion and neuronal projection (Supplementary File 7). Furthermore, a striking ∼60% and ∼80% of the 8 and 16-wpi decreasing genes, respectively, overlapped with each other or the terminally downregulated genes (Fig. 2c). Interestingly, the initial downregulation at the 8- and 16-week timepoints was transient, indicating that compensatory mechanisms might act during the earlier stages of prion disease.

To validate these changes, we analyzed the 8 wpi and terminal time point in a second, validation cohort of mice, which were inoculated and analyzed independently from the first cohort (Supplementary Fig. 3a,b and 4). Consistently, we observed that cluster 4 genes (oscillating genes) decreased at 8 wpi and the terminal stage, and that cluster 3 genes (mainly neuronal genes) decreased at the terminal stage. Furthermore, genes belonging to clusters 1 and 2 increased at the terminal stage (Supplementary Fig. 3c-f). Collectively, our data shows that hundreds of genes are downregulated at a very early disease stage, suggesting that this early response is linked to primary pathogenic events rather than reactive changes.

**Fig. 3:**
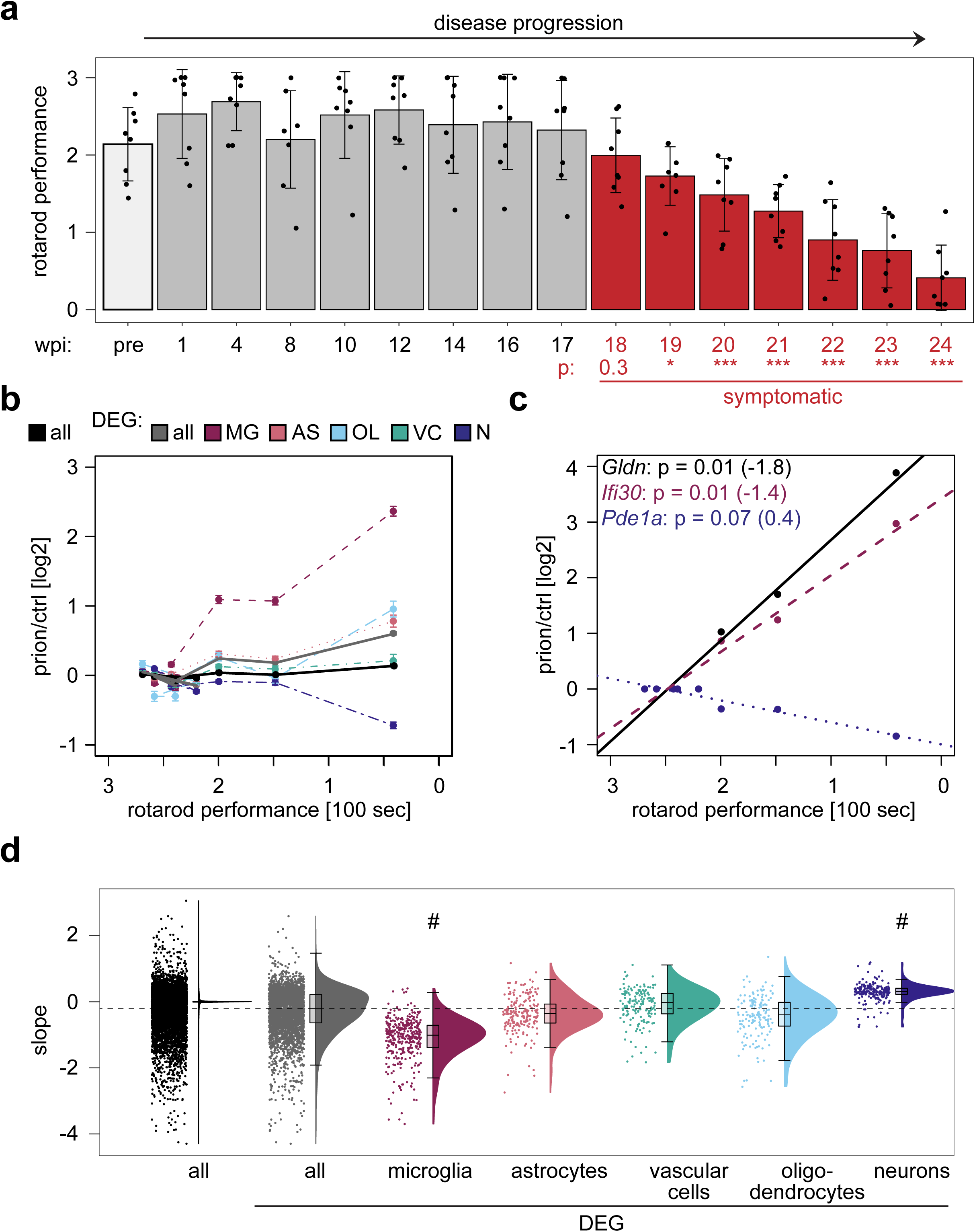
Microglia activation and prion disease progression. **a**, Rotarod performance of prion-inoculated mice during disease progression (pre = pre-inoculation). Bars represent the mean latency to fall in 100 seconds. Each dot represents one individual mouse. P-adjusted: P values were calculated with a one-way ANOVA followed by Tukey’s multiple comparison test (*p<0.05; ***p<0.001; compared to 1 wpi). Red: symptomatic phase of disease. Error bars: standard deviation. **b**, Comparison of rotarod performance (latency to fall in 100 seconds) and average log_2_FC at the corresponding time point. Lines represent all genes, all DEGs or DEGs that are enriched in the indicated cell type. Error bars: standard error of the means (SEM). **c**, Linear regression analysis between the rotarod performance and the gene expression changes of individual genes. Shown are the most significantly correlating genes that were non-enriched (black), microglia-enriched (red-violet) and neuronal-enriched (violet-blue). Corresponding Bonferroni-corrected p values and slopes of the linear regression analysis are indicated in brackets. **d**, Raincloud plot displaying the slope distribution of the linear regression analysis. The slope between all DEGs and DEGs that are either microglia- or neuron-enriched differs significantly (p < 1*10^-14^; one-way ANOVA followed by Tukey’s multiple comparison test).

**Figure 4:**
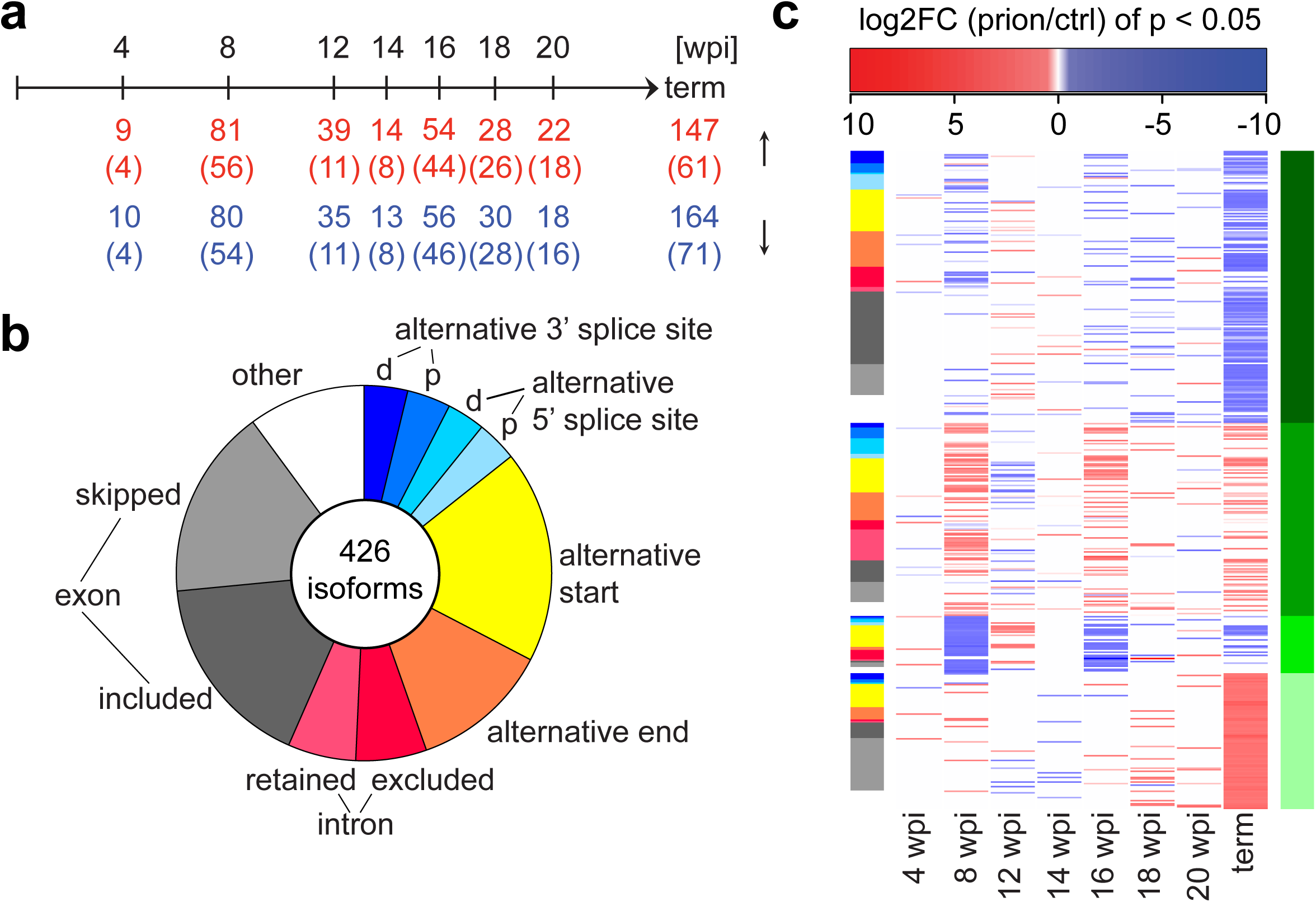
Post-transcriptional regulation during prion disease progression. **a**, Time points of prion inoculations and differentially expressed splice isoforms (FDR < 0.05 at least at one time point). The number of upregulated and downregulated isoforms (p < 0.05) at each time point and the numbers of shared genes (in brackets) are indicated. **b**, Pie chart representing splicing event percentages of 426 differentially expressed isoforms (FDR < 0.05 at least at one time point). d=distal, p=proximal. **c**, Heat map depicting the log_2_FC of differentially expressed isoforms (only log_2_FC with p < 0.05 are colored). The corresponding splicing event (same coloring as in **b**,) is indicated as a side bar on the left. Unsupervised kmeans clustering (k = 4 clusters) identified coherent patterns of log_2_FC. Clusters c1-4 are indicated as a side bar on the right.

### Microglia activation drives symptomatic prion disease progression

We next aimed at correlating the gene expression changes with the progression of neuropathological changes. Mice reached the terminal stage of the disease at 170-180 days, at which point they were sacrificed (median survival: 176 days; Supplementary Fig. 5a). Brain sections of prion-inoculated mice were assessed for morphological changes (haematoxylin/eosin), for astro- and microgliosis (GFAP and IBA1 staining) and for the presence of protease-resistant prion protein (PrP^Sc^; protease treatment followed by SAF84 staining). Spongiosis, astro- and microgliosis became evident at 16 wpi and increased at the terminal stage, whereas PrP^Sc^ started to accumulate already at 12 wpi (Supplementary Fig. 5b). These results are in line with the transcriptomic data discussed above.

We next determined the concentration of prion propagons using a quaking assay. We found that the lag phase of the assay, which measures templated nucleation of PrP fibrils, continually decreased during disease progression (Supplementary Fig. 6a), suggesting that the amounts of infectious units increased up to the terminal stage. In parallel, we analyzed prion infectivity using the scrapie cell assay in endpoint dilution format, which yielded similar results as the quaking assay and confirmed that infectivity continuously increased from 8 wpi to the terminal stage. Prion titers were significantly higher at the terminal stage compared to 16 wpi (Supplementary Fig. 6b), which is in contrast to previous reports stating that infectivity would reach a plateau at 16 wpi^7^.

To relate gene expression changes to neurological dysfunction, we assessed the motor performance of prion-inoculated and control mice using a rotarod test. We observed a progressive decline in motor performance starting at 18 wpi (Fig. 3a). The onset of impairment was synchronous to increased expression of microglia-enriched genes and was unlinked to decreased expression of neuronal genes which became evident only at the terminal stage (Fig. 3b). A linear regression analysis between rotarod performance and the expression change of each of the 3723 DEGs, identified a significant correlation for 347 DEGs (p < 0.001). Examples of correlated non-enriched, microglia-enriched and neuronal-enriched genes are shown (Fig. 3c). While the linear regression slope of microglia genes was negative, neuronal genes exhibited a positive linear regression slope. This was confirmed when we compared the average slope between DEGs enriched in different cell types (Fig. 3d). We conclude that motor decline, neuropathological changes and an increase in microglia gene expression occur simultaneously, long before terminal changes in neuronal gene expression and neuronal loss become evident. This suggests that microglia, rather than neurons, are the final drivers of prion disease progression. However, the transcriptional changes observed at 8 wpi precede both the onset of clinical signs and any changes in microglial gene expression. The early 8 wpi changes might thus hierarchically control microglia changes and ultimately induce pathogenesis.

### Identification of post-transcriptional changes during prion disease progression

Many RNA binding proteins are exclusively expressed in neurons, and aberrant splicing has been linked to multiple neurodegenerative diseases^16^. We identified a total of 426 isoforms that were differentially expressed in prion-inoculated mice at one or more time points (FDR < 0.05; Fig. 4a; Supplementary File 8). 102 of these splice isoforms mapped to DEGs, indicating that differential splicing of these transcripts might impact their abundance (Supplementary File 8). The 426 isoforms correspond to 239 splicing events, which mapped to 228 genes. Most of these genes showed just one significantly changing splicing event, except *App*, *Chl1*, *Evl*, *Fus*, *Neo1*, *Olfm1*, *Picalm*, *Ppfia4* and *Sorbs1*. The majority of splicing changes consisted of exon inclusion/skipping followed by alternative transcript starts/ends (Fig. 4b). And while most splicing changes occurred only at the terminal stage, many splicing events showed a similar trend (p < 0.05) at multiple time points during disease progression. Most strikingly, we observed an oscillating pattern of isoform expression at 8 wpi, 16 wpi and the terminal stage (Fig. 4c), indicating that these timepoints are marked by a characteristic RNA expression and splicing signature.

Select differentially spliced isoforms are shown in Supplementary Fig. 7 (indicated in Supplementary File 8). Skipping of exon 17 of *Picalm*, a susceptibility gene for late-onset Alzheimer’s disease^17^ was increased at the terminal stage of prion disease. Skipping of exon 17 introduces a premature stop codon, which leads to the production of a truncated protein, and is thought to affect clathrin-mediated synaptic endocytosis^18, 19^. The inclusion of exons 7 and 8 of *App*, a well-described splicing event^20^ increases both at 8 wpi as well as the terminal stage. Similarly, alternative splicing of the synapsin genes, *Syn1* and Syn2, and of *Ctsa*, a disease associated microglia (DAM) gene^21^ changes at both 8wpi and the terminal stage. While mRNA levels of *Ctsa* increased at the terminal stage, they remained unchanged at 8 wpi, indicating that the regulation of splicing and mRNA abundance of this gene are unlinked.

A further post-transcriptional mechanism that is essential for nervous system homeostasis is RNA editing, whose dysregulation has been implicated in neurodegeneration^22^. The most common form of RNA editing, adenosine (A) to inosine (I) conversion, is mediated by the adenosine deaminase acting on RNA (ADAR). The expression levels of the ADAR enzymes, Adar1 and Adarb2 did not change during prion disease progression, whereas Adarb1 was significantly downregulated at the terminal stage (Supplementary File 3). We did not observe major editing changes at any of the analyzed time points, even upon merging main and validation datasets (Supplementary File 9). Only two analyzed sites were differentially edited in terminally diseased mice (Supplementary Fig. 8). The two sites are in the 3’ untranslated regions of *Sh2d5* and *Padi2*, and while the mRNA expression of *Sh2d5* at that timepoint did not change, *Padi2* expression increased in terminally diseased mice.

### The impact of aging on prion disease progression

The prevalence of neurodegenerative diseases increases drastically with age. Sporadic CJD typically manifests in 55-65 years old individuals^23^. This age in humans is comparable to approximately 12 months of age in mice^24^. To assess if age-related changes accelerate prion disease progression, we compared disease progression in young vs. aged mice. We inoculated a third cohort of 2-month old “young” mice (similar to cohorts #1/2) and 1-year old mice. We analyzed gene expression changes at 8 wpi in aged mice and at the terminal stage in young and aged mice. Similar to the previously analyzed samples, we observed a separation of control and prion-inoculated samples at 8 wpi and the terminal stage (Supplementary Fig. 4 and 9a,b; Supplementary File 10). Remarkably, the median lifespan of aged prion-inoculated mice was only slightly, albeit significantly (p = 0.0001), shorter than that of young inoculated mice (Fig. 5a). This suggests that age does not strongly accelerate prion disease progression and that disease manifestation is similar in young and aged mice. Consistently, a correlation of gene expression changes was noticeable at 8 wpi (R = 0.39; Fig. 5b). Unsurprisingly, the changes in gene expression in young and aged mice converged towards the terminal stage (R = 0.89; Fig. 5c).

**Figure 5:**
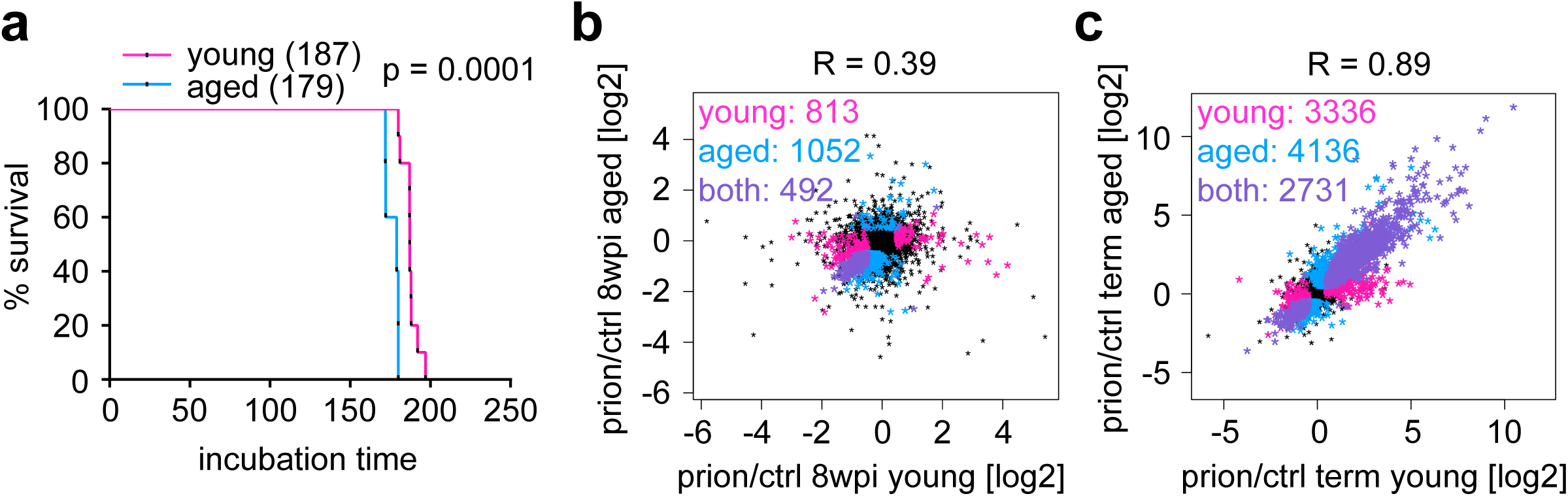
Impact of aging on prion disease progression. **a**, Survival curves of mice inoculated at 2 months (young) or one year (aged). Shown is the % survival compared to days post inoculation (dpi). The median survival, which is significantly reduced in aged mice (p = 0.0001; log-rank Mantel-Cox test) is indicated. **b**, Scatter plot comparing the log_2_FC of expressed genes (n = 15,859) in young and aged prion-inoculated mice at 8 wpi. Genes that are differentially expressed (|log_2_FC| > 0.5 and FDR < 0.05) in young (n = 813) or aged (n = 1,052) mice are colored in pink and blue, respectively. The change in expression correlates (R = 0.39) and 492 genes change both in young and aged mice (colored in purple).**c**, Scatter plot comparing the log_2_FC of expressed genes (n = 15,859) in young and aged prion-inoculated mice at the terminal stage. Genes that are differentially expressed (|log_2_FC| > 0.5 and FDR < 0.05) in young (n = 3,336) or aged (n = 4,136) mice are colored in pink and blue, respectively. The change in expression is highly correlated (R = 0.89) and 2,731 genes change both in young and aged mice (colored in purple).

### Young plasma treatment ameliorates prion disease progression

Infusion of plasma from young mice can revert age-related impairments of cognition and synaptic plasticity^25^ and ameliorates neuronal hippocampal dysfunctions in murine models of Alzheimer’s Disease^26^. We therefore analyzed the impact of young plasma on the course of prion disease as well as on the early prion-induced changes at 8wpi. Mice inoculated with prions or control brain homogenate were treated for 8 weeks with young plasma or saline via bi-weekly intravenous injections. Hippocampal samples were subjected to high-throughput sequencing at 8 wpi and at the terminal stage (Supplementary Fig. 4 and 10a,b; Supplementary File 11). We also monitored the rotarod performance and assessed mice for neurological symptoms from 18 wpi onwards by scoring for the absence/presence of a hunched posture, rigid tail, piloerection, hind limb clasping and ataxia (Fig. 6a). Remarkably, plasma administration reduced essentially all prion-induced changes at 8 wpi (Fig. 6b). Neurological deficits, which usually occur at 20 wpi, became evident only at 23 wpi in plasma-treated animals (Fig. 6c). Similarly, rotarod performance decreased at a later stage in plasma-treated samples (Supplementary Fig. 10c). In contrast, we only observed a mild, non-significant increase in the median lifespan upon plasma administration (6d), and, as expected, the terminal prion-induced changes were very similar between plasma and saline-treated mice (Fig. 6e). This suggests that young plasma treatment might improve the health span but not the lifespan of prion diseased mice.

**Figure 6:**
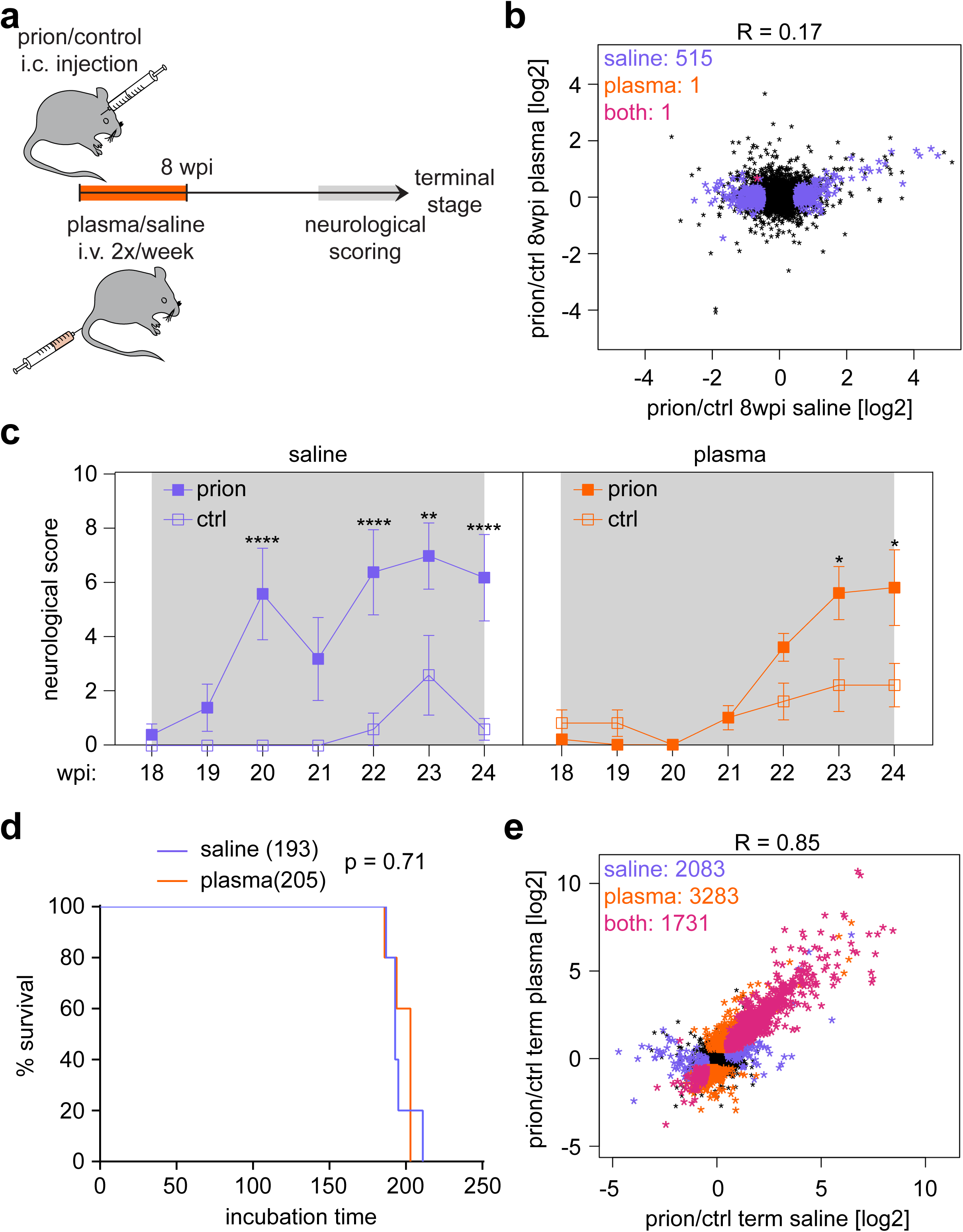
Young plasma treatment ameliorates prion disease progression. **a**, Time points of prion inoculation, plasma treatment, neurological scoring and sample analysis. **b**, Scatter plot comparing the log_2_FC of expressed genes (n = 15,863) in plasma and saline treated prion-inoculated mice at 8 wpi. Differentially expressed genes (|log_2_FC| > 0.5 and FDR < 0.05) are colored in purple (saline; n = 515) and orange (plasma; n = 1). **c**, Plot displaying the neurological score of saline and plasma-treated mice from 18 wpi until the terminal stage. Shown is the mean neurological score +/-SEM (**p<0.01; ***p<0.001; ****p<0.001; two-way ANOVA followed by Tukey’s multiple comparison test comparing prion vs control samples at each time point). **d**, Survival curves of mice that were treated for 8 weeks with saline or plasma after prion inoculation. Shown is the % survival compared to days post inoculation (dpi). The median survival of plasma treated mice did not change (p = 0.71; log-rank Mantel-Cox test). **e**, Scatter plot comparing the log_2_FC of expressed genes (n = 15,863) in plasma and saline treated prion-inoculated mice at the terminal stage. Differentially expressed genes (|log_2_FC| > 0.5 and FDR < 0.05) are colored in purple (saline; n = 2,083), orange (plasma; n = 3,283) or pink (both; n = 1,730).

## Discussion

Here we have correlated the temporal sequence of transcriptional, splicing, and RNA editing events with neuropathological changes, prion infectivity, and clinical symptoms, and have generated a searchable genome-wide database of prion-induced changes during disease progression. As expected, the histological alterations (microgliosis and astrogliosis, spongiform changes and PrP^Sc^ deposition) and clinical signs were mirrored by pronounced gene expression changes at terminal disease. Surprisingly, onset and progression of motor dysfunctions correlated precisely with the onset of glial changes and occurred long before neuronal loss was detectable. This suggests that glial perturbation, rather than neuronal demise, is the driver of prion disease progression. Moreover, we observed pronounced gene expression changes at 8 wpi, when neither neuropathological changes, prion infectivity, nor clinical symptoms are detectable. These early gene expression changes might provide new starting points for the development of novel therapeutics and diagnostics.

We further detected the differential expression of 426 splicing isoforms. Most splicing differences occurred both at terminal disease and at the preclinical 8 wpi timepoint. More than 100 of these splicing isoforms mapped to genes that were also differentially expressed, indicating that splicing and expression of these genes is linked. Nine genes were differentially spliced at multiple sites, including *App*, *Chl1*, *Evl*, *Fus*, *Neo1*, *Olfm1*, *Picalm*, *Ppfia4* and *Sorbs1*. Several of these genes have been linked to neurodegeneration, suggesting that splicing dysregulation of these genes might be common to multiple neurodegenerative diseases. In contrast, only two previously identified editing sites in 3’UTRs showed a significant change in our dataset, at the terminal disease stage. These results were inconsistent with a recent study^13^, identifying a prion-induced editing signature in mouse cortex, which could at least partially be confirmed in human autopsy brain. It remains to be seen if prion-induced editing changes differ between brain regions, or if distinct murine prion disease models and different editing analyses account for this discrepancy.

Our experiments also show that prion titers never reach a plateau but continue to rise until the terminal stage of disease. Gene expression changes occur in parallel with initial prion replication, while amounts of infectious units progressively increases over disease incubation time. Earlier investigations^6–8^ have hypothesized that scrapie neurotoxicity is only triggered when a “lethal” form of prion protein (denoted “PrP^L^”) accumulates in late pathogenesis, in a second phase of the disease, and drives toxicity and clinical disease. Conversely, prion replication would initially occur without associated toxicity. However, no physical evidence of “PrP^L^” has ever surfaced. In addition, the two-phase hypothesis rests on the quantification of prion infectivity and histological analyses fraught with analytical imprecision and poor sensitivity. The discovery of early, specific, robust and reproducible disease-associated gene expression changes before the saturation of prion infectivity refutes the two-phase hypothesis and suggests that pathology starts as soon as prions enter the brain.

Surprisingly, our analysis indicates that early alterations can be initially overcome. The existence of compensatory mechanisms is suggested by the oscillating gene expression patterns that were observed. The early downregulation of cluster-4 genes was transient and underwent complete recovery. Another peculiar expression profile was observable for certain microglial markers which were monotonically upregulated from 16 wpi. This included phagocytosis-related genes (*Aif1*, *Dock2*, *Fgr*, *Fcgr1*, *Fcgr2b*, and *Fcgr3*) and disease-associated microglia (DAM) genes (*Itgax*, *Clec7a*, *Cxcl10* and *Lag3)*^27^. Remarkably, the upregulation of these genes correlates precisely with the onset of motor deficits whereas no such correlation existed with neuronal genes. We therefore hypothesize that it is glial rather than any neuronal alterations that cause clinical disease manifestation. This aligns with gliosis being observed in symptomatic but not in pre-symptomatic CJD patients^28^. Alternatively, the dysfunction of a small number of neurons at 16-18 wpi may drive clinical signs while remaining below the detection threshold of RNAseq.

Clinically overt Creutzfeldt-Jakob disease progresses much more quickly than most other neurodegenerative syndromes, making the development of effective therapies extremely difficult. However, our data suggest that the critical period for potential intervention may be much earlier than previously estimated^6, 10^. Accordingly, infusion of young plasma during the first 8 weeks after prion inoculation attenuated the early downregulation of genes involved in neuronal functions, and also delayed the manifestation of neurological signs as observed in models of Alzheimer’s disease^26^. Interestingly, the beneficial effects of young plasma did not translate into a significantly extended overall survival, perhaps because primary prion replication was not modified despite attenuated downstream neurotoxicity. Similarly, the survival curves of young and middle-aged prion inoculated mice were similar, suggesting that ageing has only minor effects on early and late transcriptional changes as well as lifespan. And yet the plasma experiment suggests that treatment aimed at rescuing the molecular alterations identified in the clinically silent phase might impact the consequent entire progression of the disease. This idea is also in line with emerging results of novel potential therapies which seem to be effective only when started either prophylactically or within the first 11 weeks after prion inoculation (Vallabh S., et al., from Minikel E. cureffi.com; May 23, 2019). The fact that young plasma seems to improve the health span, rather than the lifespan, of prion disease may be relevant to improving the life quality of patients with neurodegenerative diseases.

It remains to be seen whether the molecular phenomena and kinetics detected in mouse models of prion disease reflect those of the corresponding human disorders. If so, the results detailed here may instruct the development of early diagnostic and therapeutic approaches. Finally, the molecular mechanisms underlying prion diseases may also be of relevance for other common protein misfolding disorders, such as Alzheimer’s and Parkinson’s disease.

## Methods

### Mice

Animal experiments were performed in compliance with the Swiss Animal Protection Law, under the approval of the Veterinary office of the Canton Zurich (animal permits ZH41/2012, ZH90/2013, ZH040/15, ZH243/15). All efforts were made to prevent or minimize animal discomfort and suffering; individual housing was avoided. Prion-inoculated and control-injected mice were regularly monitored for the development of clinical signs, according to well established procedures. Humane termination criteria were employed. Intracerebral injections, blood collection and transcardial perfusions were performed in deeply anesthetized mice. Habituation periods before experiment begin were included. Housing conditions and details of animal experimentation are detailed in the corresponding sections. C57BL/6 male mice were purchased from Charles River, Germany. Mice were kept in a conventional hygienic grade facility, constantly monitored by a sentinel program aimed at screening the presence of all bacterial, parasitic, and viral pathogens listed in the Federation of European Laboratory Animal Associations (FELASA). Animal facility was considered positive for *Murines Norovirus* and *Helicobacter spp*. Animals were kept in IVC type II long cages with autoclaved dust-free Lignocel SELECT Premium Hygiene Einstreu (140-160g/cage) (Hersteller J. Rettenmaier & Söhne GmbH), autoclaved 20 x 21 cm paper tissues (Zellstoff), autoclaved hay and a carton refuge mouse hut as nesting material. Up to five mice were housed in the same cage. Individual housing was avoided. For the plasma treatment experiment, 20 male mice were kept in groups of 5 in individually ventilated type II cages in a highly hygienic grade facility. Mice had unrestricted access to sterilized drinking water and were fed a pelleted mouse diet (Kliba No. 3340 or 3341, Provimi Kliba, Kaiseraugst, Switzerland) *ad libitum*. The light/dark cycle consisted of 12/12 h with artificial light (40 Lux in the cage) from 07:00 to 19:00h. The temperature in the room was 21±1°C, with a relative humidity of 50±5%. The air pressure was controlled at 50 Pa, with 15 complete changes of filtered air per hour (HEPA H 14 filter; Vokes-Air, Uster, Switzerland).

### Prion Inoculations

Imported mice were allowed at least one week of habituation in our animal facility before experimental manipulations. Six-week or 51-week old C57BL/6 male mice (Supplementary File 1) were injected in the right hemisphere with 30 μl of RML6 (passage 6 of Rocky Mountain Laboratory strain mouse-adapted scrapie prions) at a 10^-2^ dilution of a 10% homogenate, containing 9.02 log LD_50_ of infectious units per ml in 10% homogenate. Control inoculations were performed using 30 µl of non-infectious brain homogenate (NBH) from CD-1 mice at the same dilution. Each experimental mouse was randomly assigned a code and, based on this, to an experimental group/time point. After inoculation, mice were initially monitored three times per week. After clinical onset, mice were daily monitored. Mice were sacrificed at pre-defined time points based on their experimental group, starting always at the same time of the day and alternating between prion-inoculated and NBH-injected mice. Prion-inoculated mice allocated to the terminal group were sacrificed upon clear signs of terminal prion disease. Control-injected mice assigned to the latest time point group were sacrificed at 192 days post-inoculation, 13 days later than the last terminal prion-inoculated mouse. Transcardial perfusion with ice-cold PBS was performed in deeply anesthetized mice.

For RNAsequencing, hippocampi from both hemispheres were dissected, snap frozen and stored at −80°C until processing (Supplementary File 1). For histologic analysis (Supplementary Fig. 5b), 15 additional C57BL/6J male mice were inoculated with RML6, and groups of 3 mice were sacrificed at the indicated time points. Dissected brains were fixed in formalin. For the rotarod analysis shown in Figure 3, 8 additional C57BL/6J male mice were inoculated with RML6.

### Plasma collection and administration, and neurological scoring

Plasma preparation and administration was performed as previously described^25^, with minor modifications. Peripheral blood was collected from 150 young (6 weeks old) mice using retro-orbital bleeding in deeply anesthetized mice who were then immediately sacrificed by decapitation. Pooled mouse plasma was separated from blood collected with EDTA by centrifugation at 1,000*g*, followed by dialysis using 3.5-kDa D-tube dialyzers (EMD Millipore) in PBS to remove EDTA, aliquoting and storage at −80 °C until use. Systemic administration of either plasma or saline as control was performed by injection of 100µl in the tail vein of mice who had received an intra-cerebral injection with either RML6 prions or NBH. Treatment started the day after the intracerebral injection and was performed twice weekly for 8 weeks. Mice were sacrificed at 8 wpi or the terminal stage (n=5 per each group) by transcardial perfusion with ice-cold PBS in deeply anesthetized mice. Hippocampi from both hemispheres were dissected, snap frozen and stored at −80°C until processing. RNA extraction and sequencing were performed in two separate rounds (Supplementary File 1) but were pooled for analysis.

Neurological scoring was performed weekly from 18 wpi until 24 wpi by an observer who was blinded to experimental group allocation (RML or NBH-injected, plasma-or saline-treatment), and encompassed the assignment of scores from 0 (no abnormality) to 1-2-3 (for mild, moderate or severe abnormalities) in the following domains: hunched posture, piloerection, rigid tail, hind limbs clasping, ataxia.

### Immunohistochemistry

Histological analyses were performed on 2-µm thick sections from formalin fixed, formic acid treated, paraffin embedded brain tissues, as previously described^29^. Sections were subjected to deparaffinization through graded alcohols, followed by heat-induced antigen retrieval performed in EDTA-based buffer CC1 (Ventana). Stainings were performed on a NEXES immunohistochemistry robot (Ventana instruments) with the following antibodies: Iba1 (1:1000, Wako); GFAP (1:13000, Dako); SAF84 (1:200, SPI bio). For the latter staining, incubation with protease 2 (Ventana) was performed before antibody incubation. Immunoreactivity was visualized using an IVIEW DAB Detection Kit (Ventana). Haematoxylin and eosin stainings were performed according to standard procedures. Slides were scanned with NanoZoomer and images were acquired using the NanoZoomer Digital Pathology System (NDPview, Hamamatsu Photonics).

### Rotarod

The rotarod test was performed as previously described^29^, with minor modifications. The rotarod apparatus (Ugo Basile) consisted of a rotating cylinder (ø 3 cm) subdivided in five 57-mm wide lanes by dividers (ø 25 cm). Each test consisted in habituation phase and an experimental phase. The habituation phase comprised three sessions of 1 min each, at a constant speed of 5 rotations per minute (rpm), with inter-sessions intervals of at least 15 min. The test phase, which started at least 15 min after the last habituation trial, comprised three sessions of maximum 5 min each, at a constant acceleration from 5 rpm to maximum of 40 rpm, with inter-sessions intervals of at least 15 min. Latency to fall was calculated as the time until either falling from the drum or clinging to the rod and passively rotating with occurred. Rotarod tests were performed at the same time of the day (between 2 pm and 4 pm).

### Standard Scrapie Cell Assay

CAD5 cells were grown with standard OFBS Medium (Opti-MEM containing 10 % FBS, 1 % streptomycin and penicillin, 1 % Glutamax; Gibco) in a T150 cell culturing flask. One day prior to infection, 10,000 CAD5 and CAD5 KO cells lacking PrP^C^ expression were plated with 100 µL OFBS in 96-well cell culture plates (TPP) and incubated at 37 °C with 5% CO_2_. On the following day, 100 µL of brain homogenate diluted in OFBS mixed with 0.01% brain homogenate from C57BL/6J-*Prnp*^ZH3/ZH3^ mice to provide a complex matrix was added to the cells for the infection. To establish a standard curve for infection, a 1:5 serial dilution of RML6 brain homogenate (20% w/v in 0.32M sucrose, 10 LD_50_ units per mL) was used with a range from 1 × 10^−3^ to 6.4 × 10^-8^. For each sample, three different dilutions were performed ranging from 1 × 10^−3^ to 1 × 10^−5^. To control for residual inoculum, CAD5 KO cells were incubated with RML brain homogenate corresponding to the highest concentration of the standard (0.01%). CAD5 cells were incubated with (0.01%) non-infectious brain homogenate (10% w/v in 0.32M sucrose) to control for efficient proteinase K (PK) (Roche) digestion and to compute the background of the assay. Three days following infection, cells were split 1:8 into new 96 well plates containing fresh OFBS. After reaching confluence, two additional 1:8 splitting steps were performed, corresponding to days 7 and 10 post infection. On day 14 post infection, ELISPOT membranes (Millipore) were activated by adding filtered 50 µL ethanol/well, washed twice with 160 µL PBS and nearly 40,000 cells per well transferred onto the membrane and dried with a plate thermomixer (Eppendorf) at 50 °C. After drying, plates were stored at 4 °C until lysis and digestion. 50 µL of 0.5 ug/mL PK in lysis buffer (50 mM Tris-HCl pH8, 150 mM NaCl, 0.5% w/v sodium deoxycholate, 0.5% w/v Triton-X-100) was added to each well and incubated for 90 minutes at 37 °C. Following incubation, vacuum was applied to discard the contents and wells were washed twice with 160 µL PBS. To stop digestion, 160 µL of 2 mM PMSF (Sigma Aldrich) diluted in PBS was applied to the membrane and incubated at room temperature for 10 min. Tris guanidinium thiocyanate was prepared by diluting 3 M guanidinium thiocyanate in 10 mM Tris HCl pH8, and added subsequently with a total volume of 160 µL/well and incubated for 10 min. Supernatant was discarded into 2M NaOH and membrane was washed seven times with each 160 µL PBS and blocked 1 h with 160 µL Superblock (Thermo Scientific) prepared in MilliQ. Remaining blocking solution was removed under vacuum and 50 µL POM1 antibody was applied at a concentration of 1:5000 diluted in TBST (10 mM Tris HCl, pH 8, 150 mM NaCl, 0.1% (v/v) Tween 20) containing 1% (w/v) non-fat dry milk for 1 h. Supernatant was discarded and wells were subsequently washed seven times with TBST under vacuum. 50 µL of anti-IgG1-AP (Southern Biotechnology Associates) was used with a 1:4500 dilution in TBST-1% (w/v) non-fat dry milk and incubated for 1 h. Discarding of the supernatant and washing was performed in the same way as for the POM1 antibody. 50 µL of AP dye (Bio-rad) for the reaction was applied and incubated for 16 minutes. Membrane was washed twice with water, dried and stored at −20 °C in dark. Membranes were quantified using ImageJ (open source) with optical density, to distinguish between spots (representing cells that contain PK-resistant PrP) and clear areas. Data derived from SSCA have also been used for a scaling analysis to estimate rates of replication *in vivo* (Meisl et al., submitted).

### Real-time quaking induced conversion assay (RT-QuIC)

RT-QuIC assays of prion-infected mouse brain homogenates were performed as previously described^30^. Briefly, recombinant full-length hamster PrP (23–231) was expressed in Rosetta2(DE3)pLysS *E.coli* competent cells and purified from inclusion bodies by affinity chromatography using Ni^2+^-nitrilotriacetic acid Superflow resin (QIAGEN). Recombinant hamster PrP (HaPrP) was used as monomeric substrate protein for the RT-QuIC conversion. Mouse brain homogenates (20% in 0.32M sucrose) were diluted 2000-fold and used as seeds for the RT-QuIC conversion. RT-QuIC reactions containing HaPrP substrate protein at a final concentration of 0.1 mg/ml in PBS (pH 7.4), 170 mM NaCl, 10 μM EDTA, 10 μM Thioflavin T were seeded with 2 μl of diluted brain homogenates in a total reaction volume of 100 μl. NBH- and RML6-treated brain homogenates were used as negative and positive controls, respectively. The RT-QuIC reactions were amplified at 42 °C for 100 h with intermittent shaking cycles of 90 s shaking at 900 rpm in double orbital mode and 30 s resting using a FLUOstar Omega microplate reader (BMG Labtech). Aggregate formation was followed by measuring the thioflavin T fluorescence every 15 min (450 nm excitation, 480 nm emission; bottom read mode).

### RNA, extraction, library preparation and sequencing

RNA was extracted from snap frozen brain areas (both hemispheres for hippocampus) using the RNeasy Plus Universal Kit (QIAGEN), following manufacturer’s instructions. Quantity and quality of RNA were analyzed with Qubit 1.0 Fluorometer (Life Technologies) and Bioanalyzer 2100 (Agilent Technologies), respectively. The TruSeq RNA Sample Prep kit v2 (Illumina) was employed for library preparation. In brief, 1 μg of total RNA per sample was poly A enriched, reverse transcribed into double-stranded cDNA and then ligated with TruSeq adapters. PCR was performed to selectively enrich for fragments containing TruSeq adapters at both termini. Quantity and quality of enriched libraries were analyzed using Qubit (1.0) Fluorometer and Caliper GX LabChip GX (Caliper Life Sciences). The resulting product is a smear with a mean fragment size of approximately 260 bp. Libraries were then normalized to 10 nM in Tris-Cl 10 mM, pH 8.5, with 0.1% (vol/vol) Tween 20. Cluster generation was performed with the TruSeq PE Cluster kit v4-cBot-HS (Illumina), using 2 pM of pooled normalized libraries on the cBOT. Sequencing was performed on Illumina HiSeq 4000 paired-end at 2 × 126 bp using the TruSeq SBS kit v4-HS (Illumina).

### Data analysis

#### Alignment and feature counting

RNA sequencing data analysis was performed as previously described^31^, with modifications. Quality control of reads was performed using FastQC. Low-quality ends were clipped (5’: 3 bases; 3’: 10 bases). Trimmed reads were aligned to the reference genome and transcriptome (FASTA and GTF files, respectively, downloaded from the UCSC mm10) with STAR version 2.3.0e_r291^32^ with default settings.

#### Differential gene expression

Differentially expressed genes were identified based |log_2_FC| > 0 and FDR < 0.05 using the *R* package *edgeR*^33^ from Bioconductor (version 3.0). Only genes with at least 10 counts in at least 50% of the samples in one of the groups were considered in the analysis. Differentially expressed genes (DEGs) were defined as genes changing with |log_2_FC| > 0.5 and FDR < 0.05. Log_2_FC with p ≥ 0.05 were set to 0 for unsupervised kmeans clustering, t-distributed stochastic neighbor embedding (t-SNE) and the comparison with rotarod performance (Fig. 1b, c, 3b – d, and Supplementary Fig. 2a). RNA isolated from saline and plasma-treated mice was isolated, processed and sequenced in two independent runs (indicated in Supplementary Fig. 10a, b). All saline and plasma-treated mice were analyzed together. An additional covariate was added to the model in edgeR to account for the batch effect associated with the different runs. To assess if DEGs changed at multiple time points (Fig. 2a, c and Supplementary Fig. 2b) we lowered the cut-off to |log_2_FC| > 0.5 and p < 0.05.

#### Alternative splicing

Analysis of splice variants was performed by using SGseq^34^ and DEXseq^35^ *R* packages as described previously^36^ with some modifications. SGseq was first applied to identify splicing events (*e.g.* cassette alternative exon) related to two or more variants (*e.g.* isoforms with exon included or exon skipped). Exons and splice junctions were predicted from BAM files. Predictions for each sample were merged to obtain a common set of transcript features, and exons were partitioned into disjoint exon bins. A genome-wide splice graph was assembled based on splice junctions and exon bins, and splice events were identified from the graph. To determine differential splicing events, a single value for each variant was produced by either adding up the 5’ and 3’ counts, or, if these represented the same transcript features, by considering the unique value. These counts then constituted the initial input to DEXSeq, as described in the SGSeq manual. Briefly, instead of quantifying differential usage of exons across a single gene, we analyzed differential usage of variants across a single event. Such adaptation of DEXSeq is also reported in the DEXSeq vignette. Similar to the differential gene expression analysis, we retained only variants with at least five counts in at least three samples (of any condition). After this filtering step, in events associated with a unique variant, such variant was considered to be constitutive and discarded. For most comparisons, these two filters reduced the total number of variants tested to around 6,000. Differential analysis was then performed implementing a sample+exon+condition: exon model in DEXSeq. To limit the number of tests, the first variant of each event (generally a ‘skipping’ variant) was discarded. As our data, possibly due to the Nugen amplification, shows high levels of intronic reads, retained intron events were also discarded. Differentially expressed isoforms were defined as isoforms changing with FDR < 0.05. Log_2_FC with p ≥ 0.05 were set to 0 for unsupervised kmeans clustering (Fig. 4c). To compare if splicing isoforms were differentially expressed at multiple time points (Fig. 4a) we lowered the cut-off to p < 0.05.

#### RNA editing

Examination of RNA editing was conducted as previously described^37^. A catalogue of loci with mismatches with respect to the reference genome was created following RNAsequencing based best practice workflow compiled for GATK version 3.4.^38^. 17,831 RNA editing sites have previously been reported in mice^39^, and were further investigated. For the identification of differentially edited loci, data from both cohort#1 and cohort#2 were analyzed together. For each locus we considered a summary statistic (the sum) of the counts of the allelic observations across the samples in each group. In the pair-wise comparisons, we then constructed a contingency table based on edited/unedited observations at the locus in the prion/ctrl groups. An exact Fisher test was performed, and the p-values were Benjamini-Hochberg adjusted for multiple testing.

#### Database comparisons

As previously described^37^ a list of genes enriched in neurons, astrocytes, oligodendrocytes, microglia and CNS endothelial cells (500 genes per cell type) was retrieved using the cell type enrichment query from a transcriptomic based database, (http://web.stanford.edu/group/barres_lab/brain_rnaseq.html)^15^, searching for genes enriched in one cell type respect to all others. In the case of oligodendrocytes, myelinating oligodendrocytes, newly formed oligodendrocytes and oligodendrocytes precursor cells were considered together. The resulting, non-overlapping list of CNS cell-type enriched genes was used for comparisons with the lists of genes with differential gene expression, alternative splicing and RNA editing in the present study.

Gene Ontology analyses were performed with GORILLA (http://cbl-gorilla.cs.technion.ac.il), comparing indicated gene lists with a list of expressed genes.

#### Data visualization

Data was visualized in R using the packages DESeq2, RColorBrewer, pheatmap, gplots, ggplot2, NbClust, factoextra, eulerr, tidyverse, datasets, data.table, and circlize. t-distributed Stochastic Neighbor Embedding (t-SNE) method was used for projecting high dimensional gene expression data in a 2D space. We used the t-SNE implementation from scikit-learn library in Python with the following parameters: learning rate=200, n iter=1,000, random state = 0, metric=Euclidean, init=pca.

### Data availability

All data will be deposited in the GEO database. In addition, gene expression profiles during disease development can be easily visualized and downloaded at the following website: http://histodb12.usz.ch/iMice/public/PrionRNASeqDatabase/PrionRNASeqDatabase.php

### Code availability

Code is available from the authors by request.

## Acknowledgements

The authors acknowledge Catherine Aquino Fournier for RNA sequencing; Olga Romashkina, Mirzet Delic, Ahmet Varol, Laura Cervantes, Karina Arroyo, Marianne König, Irina Abakumova, Rita Moos for excellent technical help; Magdalini Polymenidou, Marc Emmenegger, Alessandro Crimi, Assunta Senatore, Mark D. Robinson, Constance Ciaudo, Daniel Spies for discussion on data analyses, and members of the Aguzzi laboratory for critical comments, Tony Wyss-Coray and Joseph M. Castellano for sharing their protocol for young-plasma treatment and for discussions, and Simon Stephan and Norbert Wey for building the searchable database. A.A. is the recipient of an Advanced Grant of the European Research Council, the Swiss National Foundation, the Clinical Research Priority Programs “Small RNAs” and “Human Hemato-Lymphatic Diseases”, the Nomis Foundation and a donation from the estate of Dr. Hans Salvisberg. S.S. is recipient of a grant from the Synapsis foundation. M.N. is recipient of a grant from the Amyloidosis Foundation. C.S. was supported by a Marie Curie Individual Fellowship.

## Author information

Silvia Sorce and Mario Nuvolone contributed equally.

## Contributions

S.S., M.N., A.A.: conception and design of the work; S.S., M.N., D.H., M.A., M.P., S.H., P.S., M.D., C.S.: acquisition and/or analysis of data; G.R., A.C., D.S.: creation of data processing and visualization tools used in the work; S.S., M.N., C.S., A.A.: data interpretation, manuscript drafting. All authors were involved in manuscript preparation and editing, approved the final version and agreed with the submission of the work.

## Ethics declarations

All authors declare no competing interest.

## Supplementary Figure Legend

**Supplementary Figure 1:**
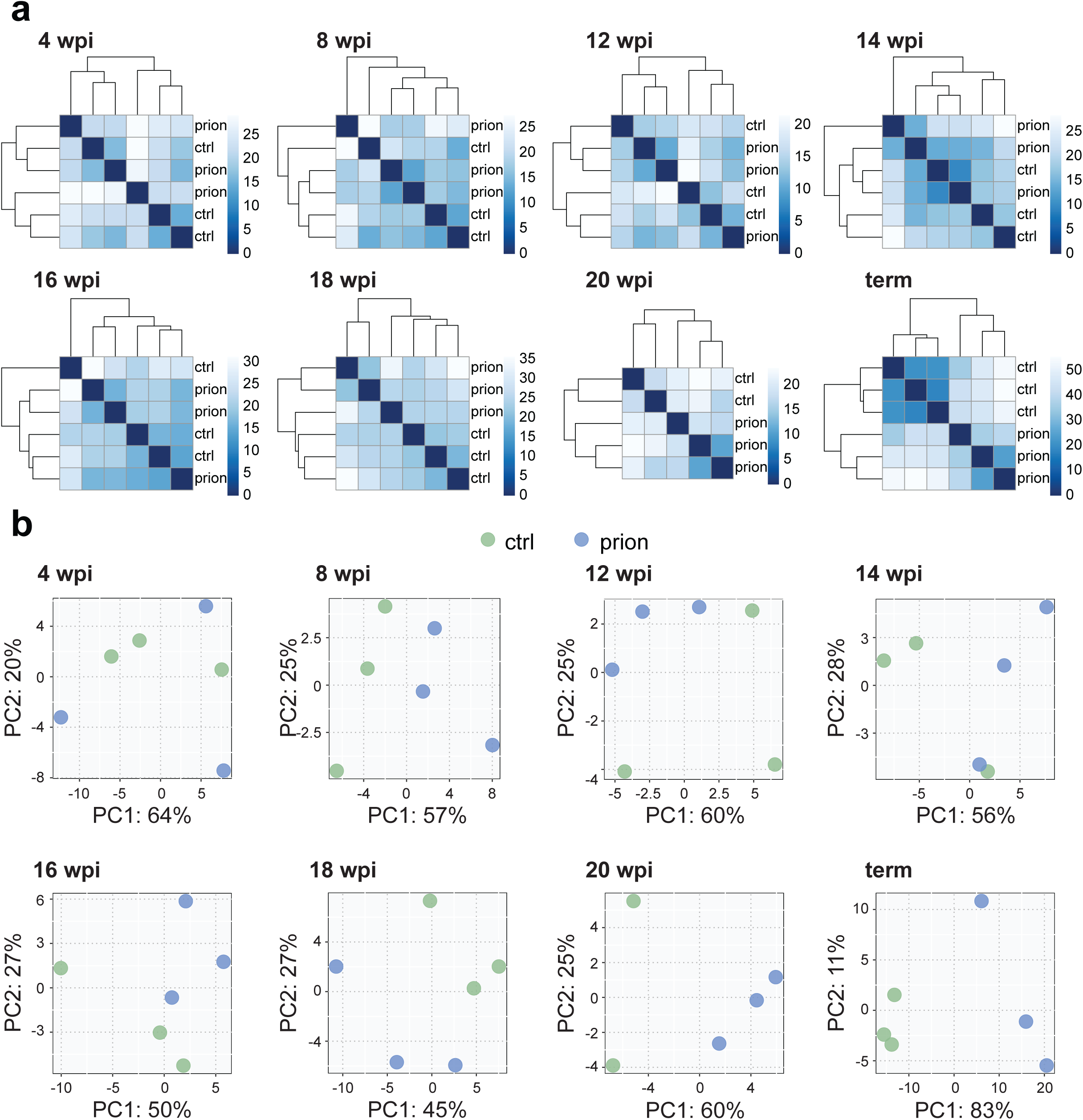
Sample separation during prion disease progression. **a**, Hierarchical clustering based on Euclidean distances. Heatmaps depicting the sample distances at each time point based on RNAseq expression data. Control and prion-injected samples cluster from 18 wpi onwards. **b**, Principal component analysis of RNAseq samples at different time points revealing a separation of control (green) and prion-injected (blue) samples at 8 wpi, 20 wpi and the terminal stage.

**Supplementary Figure 2:**
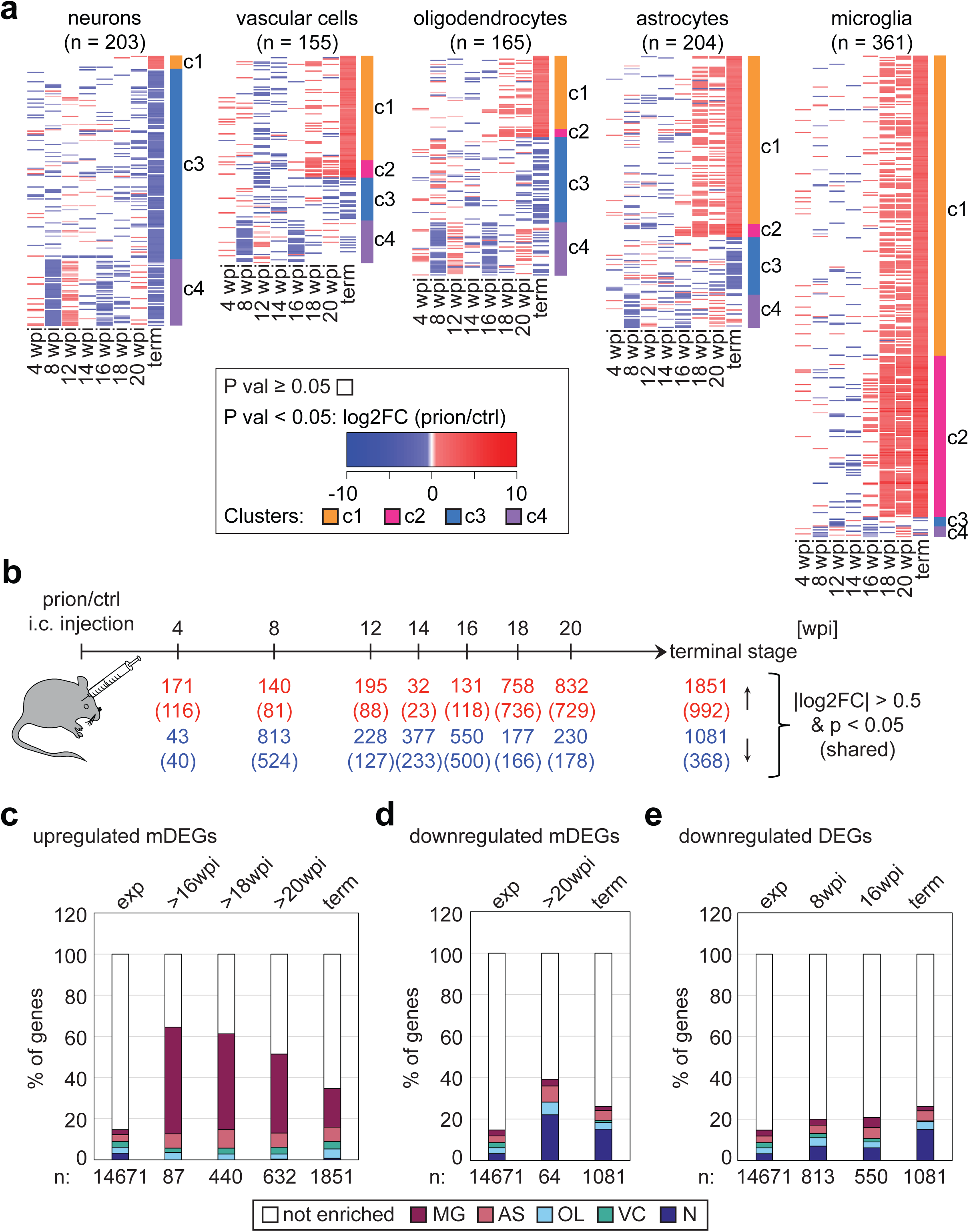
Distribution of cell-type enriched genes among DEGs. **a**, Heat maps depicting the log_2_FC of DEGs known to be enriched in different cell types (only log_2_FC with p < 0.05 are colored). The corresponding clusters are indicated as a side bar. The height of the heatmap corresponds to the number of genes. **b**, Schematic depicting time points of prion inoculations. Number of upregulated and downregulated DEGs (|log_2_FC| > 0.5 and p < 0.05), and numbers of shared genes (in brackets) are indicated. **c-e**, Percentages of cell-type enriched genes within upregulated mDEGs (c), downregulated mDEGs (d) and downregulated genes at different time points (e).

**Supplementary Figure 3:**
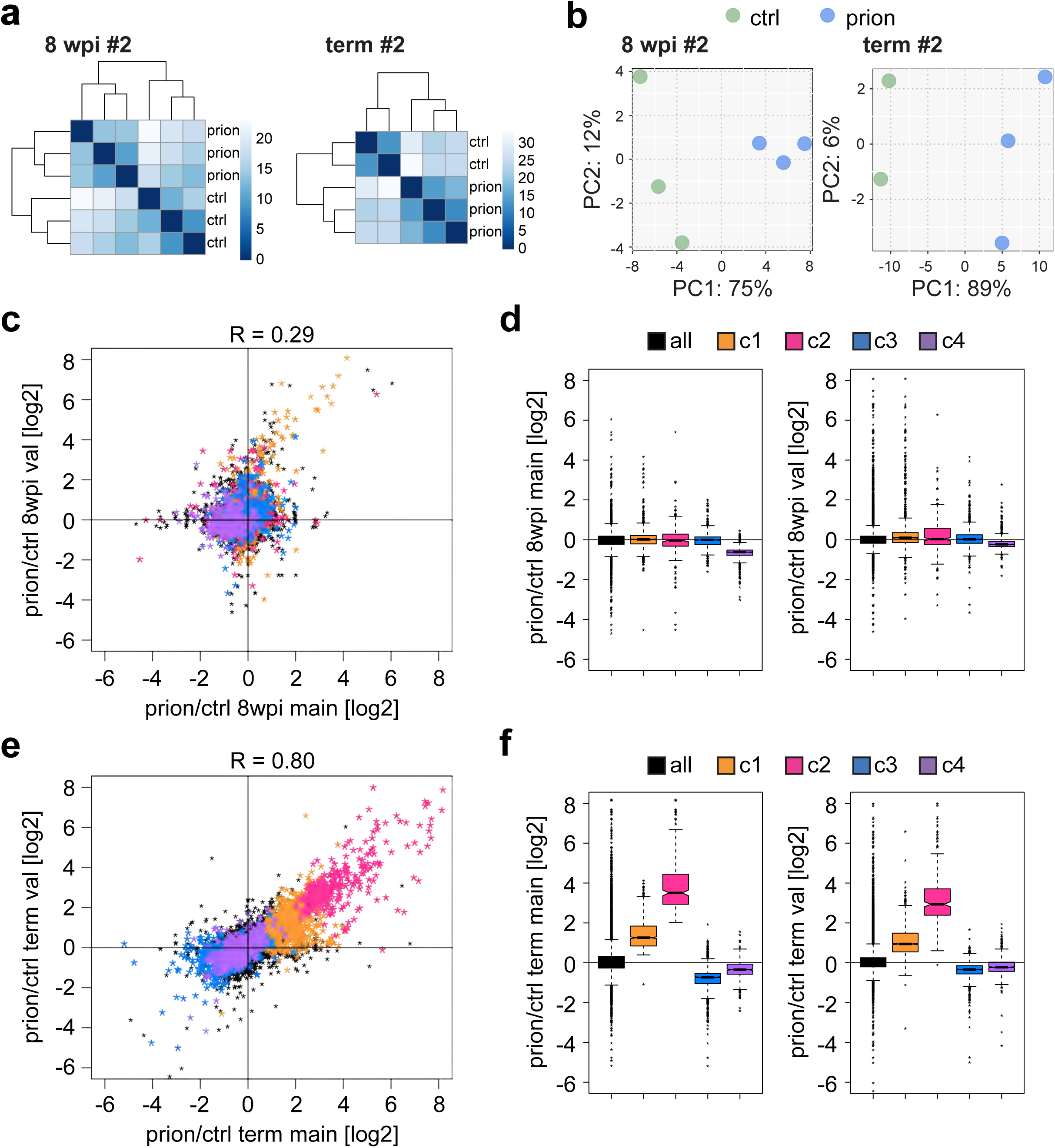
Validation of mRNA expression changes in a second, validation cohort. **a**, Hierarchical clustering based on Euclidean distances. Heatmaps depicting the sample distances based on RNAseq expression data. Control and prion-injected samples cluster at 8 wpi, and the terminal stage. **b**, Principal component analysis of RNAseq samples revealing a separation of control (green) and prion-injected (blue) samples at 8 wpi, and the terminal stage. **c**, Scatter plot depicting expressed genes (n = 15,911). The change in expression at 8 wpi correlates between the main and the validation datasets (R = 0.29). Genes belonging to the clusters identified in Fig. 1 are colored. **d**, Boxplots representing the log_2_FC distribution in the main (left panel) and validation (right panel) datasets at 8 wpi. Only expressed genes that are included (n = 15,911). The distribution of all genes (black) and genes belonging to the clusters identified in Fig. 1 are shown. **e**, Scatter plot depicting expressed genes (n = 15,911). The change in expression at the terminal stage highly correlates (R = 0.80) and genes belonging to the clusters identified in Fig. 1 are colored. **f**, Boxplots representing the log_2_FC distribution in the main (left panel) and validation (right panel) datasets at the terminal stage. Only expressed genes that are included (n = 15,911). The distribution of all genes (black) and genes belonging to the clusters identified in Fig. 1 are shown.

**Supplementary Figure 4:**
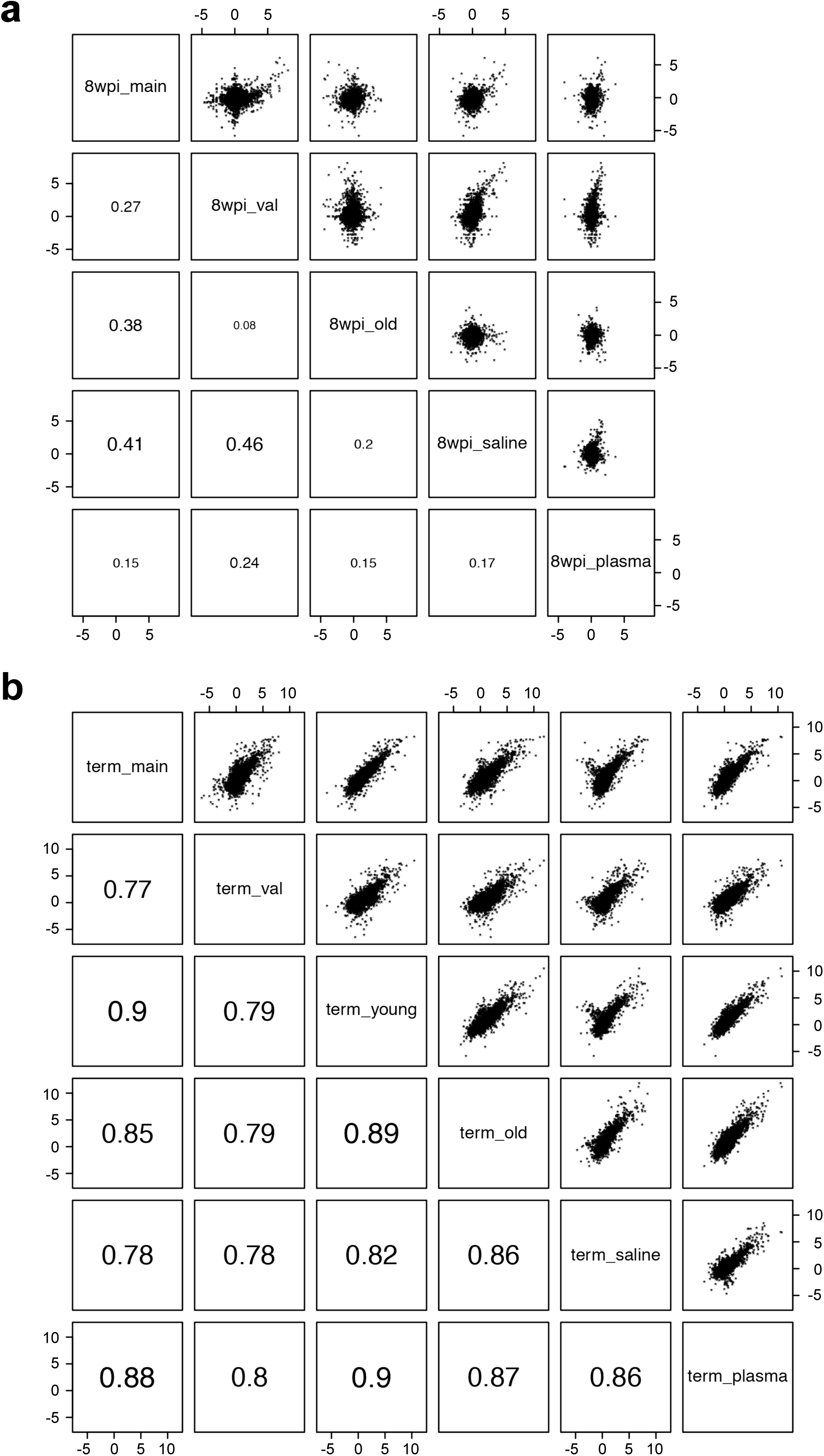
Cross-correlation plots comparing all 8 wpi and terminal samples. **a**, Cross-correlation plot of all 8 wpi comparisons included in this manuscript (main, validation, aged, saline-treated, plasma-treated). Shown are scatter plots comparing the 8 wpi log_2_FC between the groups and the corresponding R values (scaled according to R value). The plasma-treated sample correlates least with the other 8 wpi samples. Only expressed genes are included (n = 16,371). **b**, Cross-correlation plot of all terminal comparisons included in this manuscript (main, validation, young, aged, saline-treated, plasma-treated). Shown are scatter plots comparing the terminal log_2_FC between the groups and the corresponding R values (scaled according to R value). All samples strongly correlate with each other. Only expressed genes are included (n = 16,371).

**Supplementary Figure 5:**
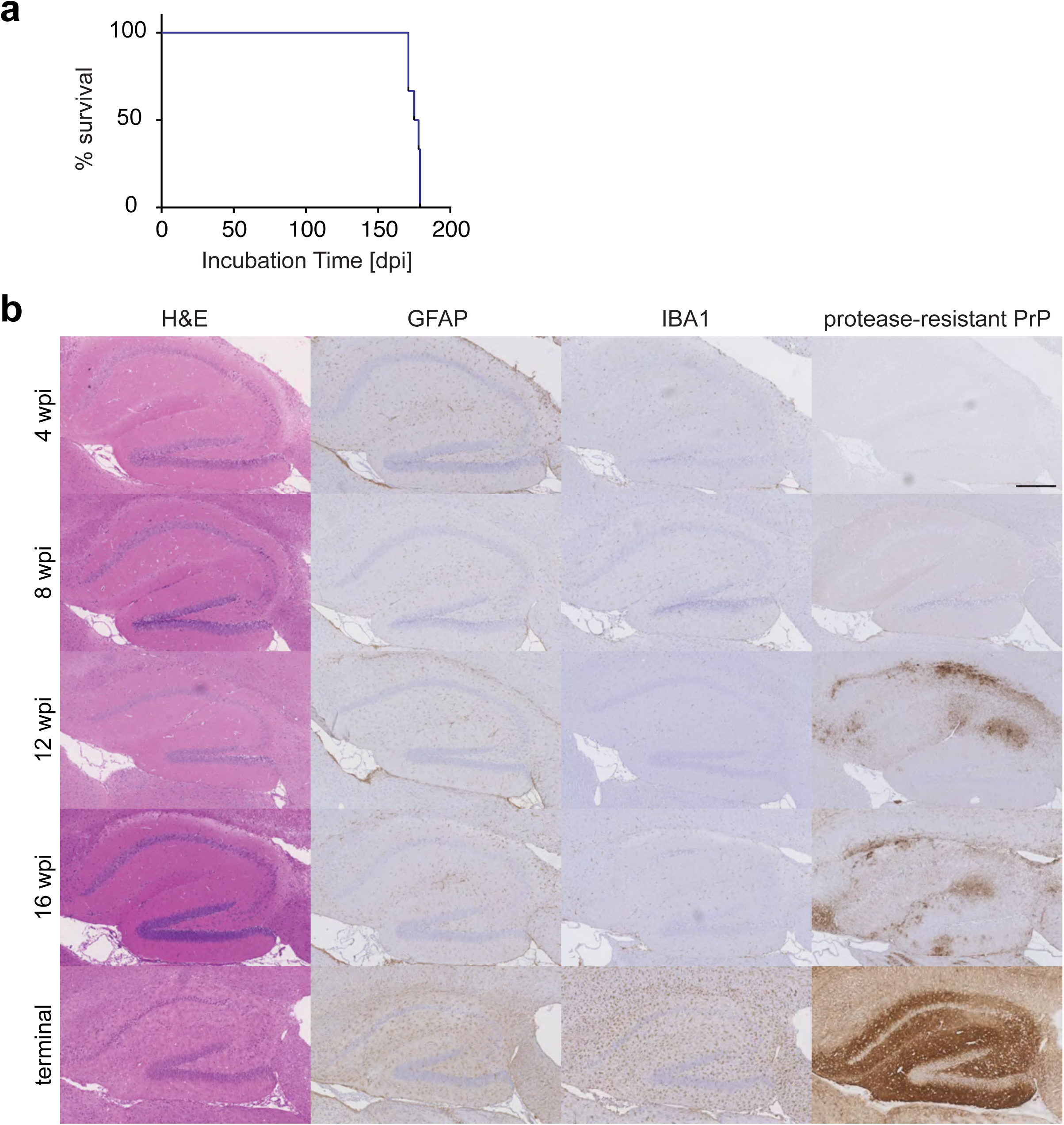
Characterization of neuropathological changes. **a**, Survival curve of mice sacrificed at the terminal stage. Shown is % survival compared to days post inoculation (dpi). **b**, Prion-inoculated mice were sacrificed at the indicated time points during disease progression. Brain section were stained with hematoxylin and eosin (H&E), GFAP (astrocyte marker), IBA1 (microglia marker) and SAF84 (detects only PrP^Sc^ after protease treatment). Scale bar in upper right panel: 250µm (applicable to all panels).

**Supplementary Figure 6:**
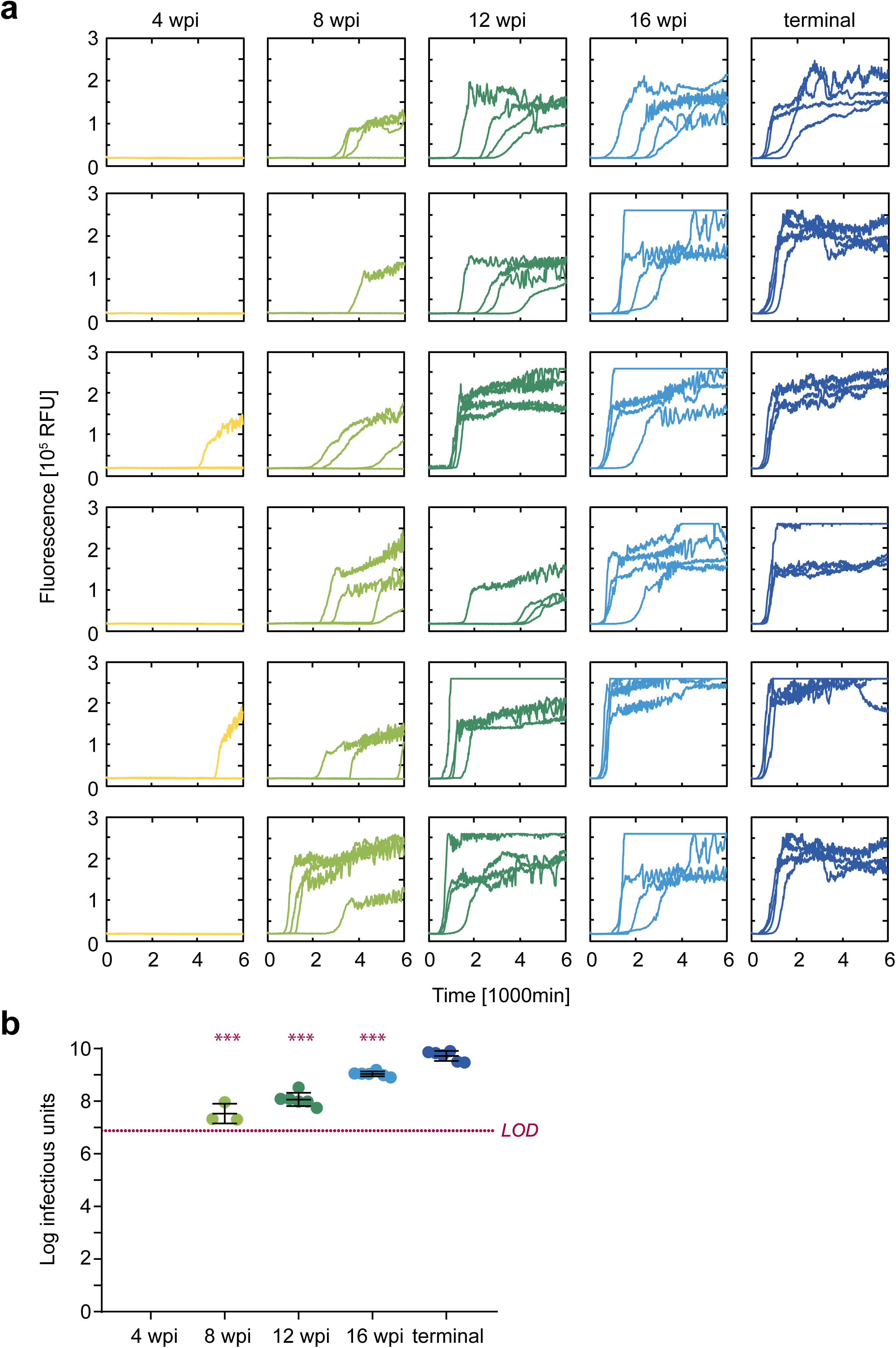
Assessment of prion infectivity. **a**, RT-QuIC reactions using brain homogenates of mice inoculated with prions and sacrificed at indicated time points. Each sample was tested in quadruplicates, and each plot corresponds to one mouse (n = 6). RFU: relative fluorescence units. **b**, Dot plot graph showing infectious units measured by standard scrapie cell assay (SSCA). Each dot represents one mouse, bars indicate standard deviations. 6 (out of 6) samples at 4wpi and 3 (out of 6) samples at 8 wpi were below the limit of detection (LOD). P values were calculated with a one-way ANOVA followed by Tukey’s multiple comparison test (***p<0.001; compared to terminal).

**Supplementary Figure 7:**
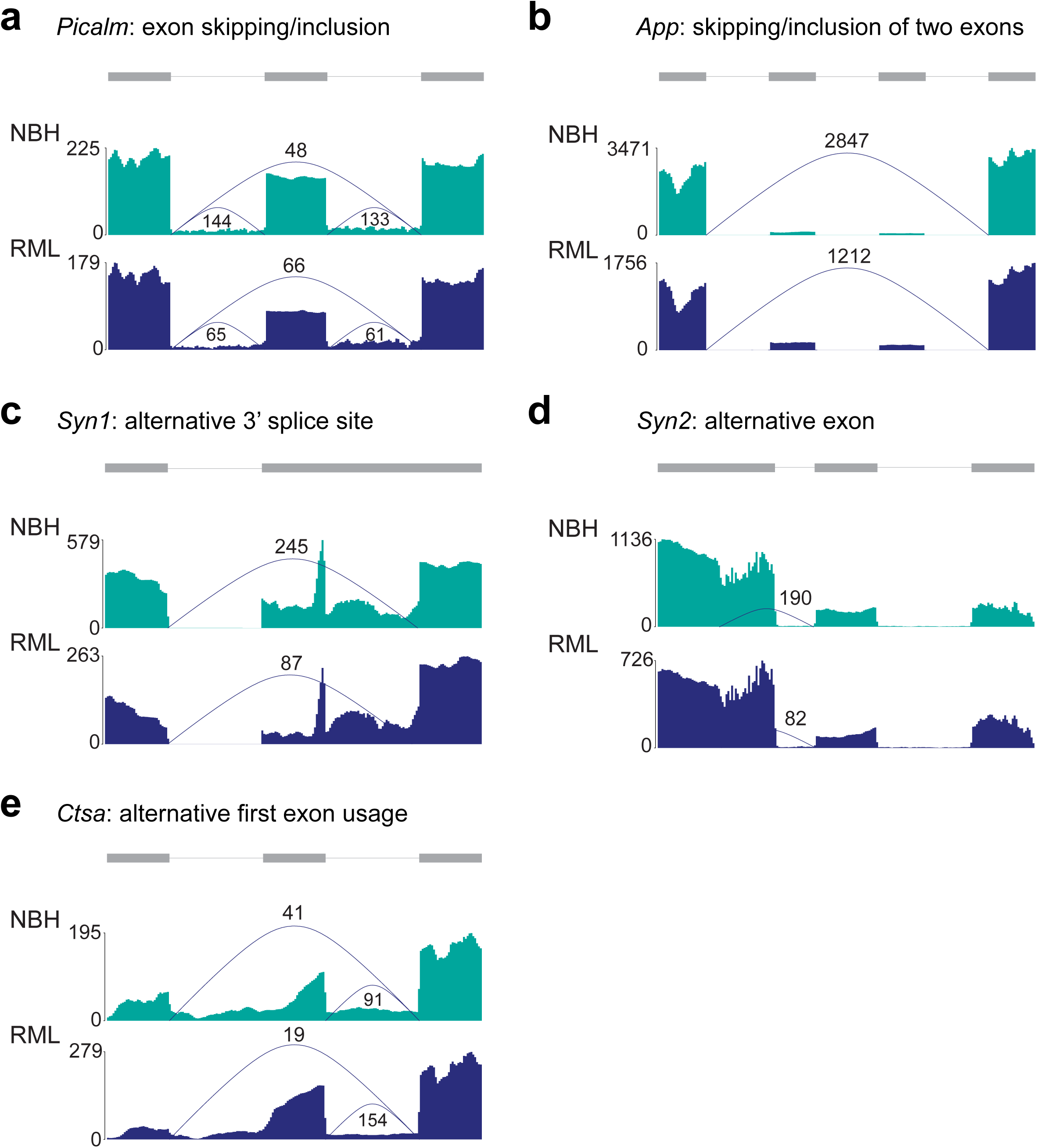
Differentially spliced isoforms during prion disease progression. (Alternative) exons are shown in grey. Exons and introns are not drawn to scale. Average per-base exon read coverage and junction counts normalized to total read counts in control and prion diseased mice at the terminal time point are shown. Splicing events were visualized with the plotSpliceGraph function of the SGSeq package in R. Shown events are indicated in Supplementary File 8.

**Supplementary Figure 8:**
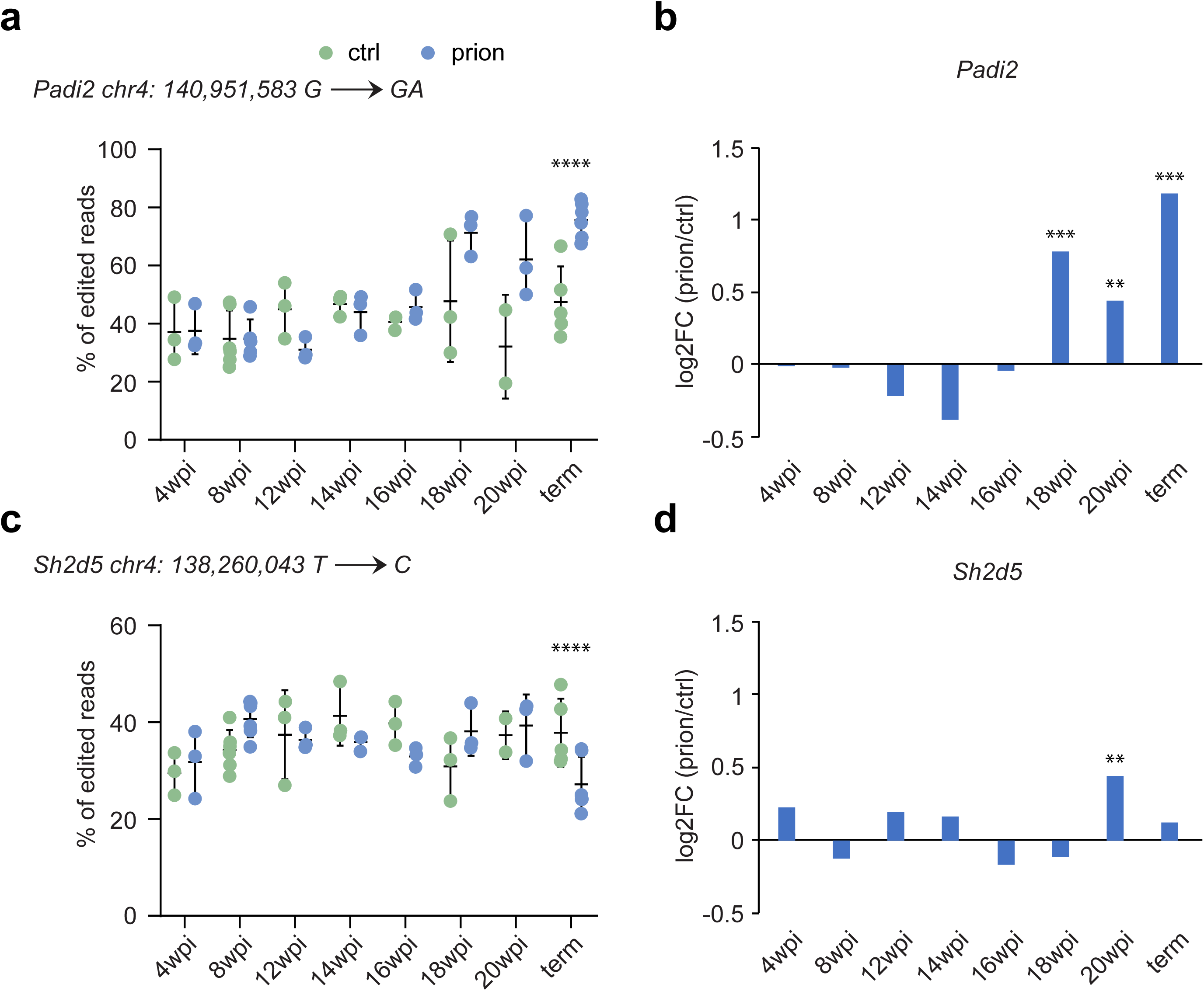
RNA editing during prion disease progression. **a and c**, Scatterplot displaying the % of edited reads (of total reads) for *Padi2* (a) and *Sh2d6* (c) in ctrl and prion diseased mice at the indicated time points. The difference in editing between ctrl and prion samples was assessed with a Benjamini-Hochberg adjusted Fisher’s exact test (****p.adj<0.0001). Error bars: standard deviation. b and d, Bargraphs representing the log2FC in mRNA expression (prion/ctrl) for *Padi2* (b) and *Sh2d6* (d) at the indicated time points. Values were derived from edgeR (**FDR<0.1; ***FDR< 0.05).

**Supplementary Figure 9:**
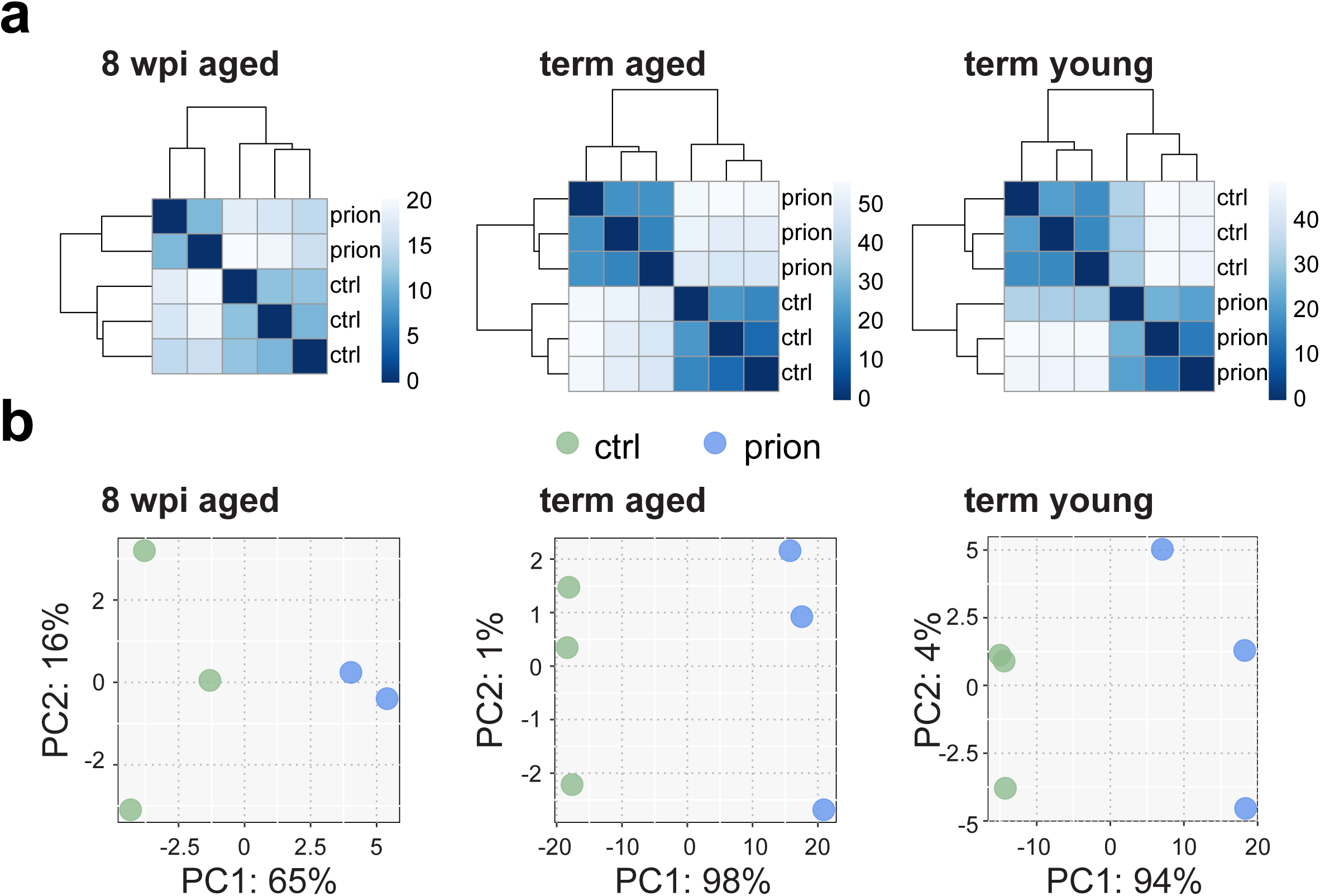
Impact of aging on prion disease progression. **a**, Hierarchical clustering based on Euclidean distances. Heatmaps depicting the sample distances based on RNAseq expression data. Control and prion-injected samples cluster at 8 wpi and the terminal stage in aged mice, and at the terminal stage in young mice. **b**, Principal component analysis of RNAseq samples revealing a separation of control (green) and prion-injected (blue) samples at 8 wpi and the terminal stage in aged mice, and at the terminal stage in young mice.

**Supplementary Figure 10:**
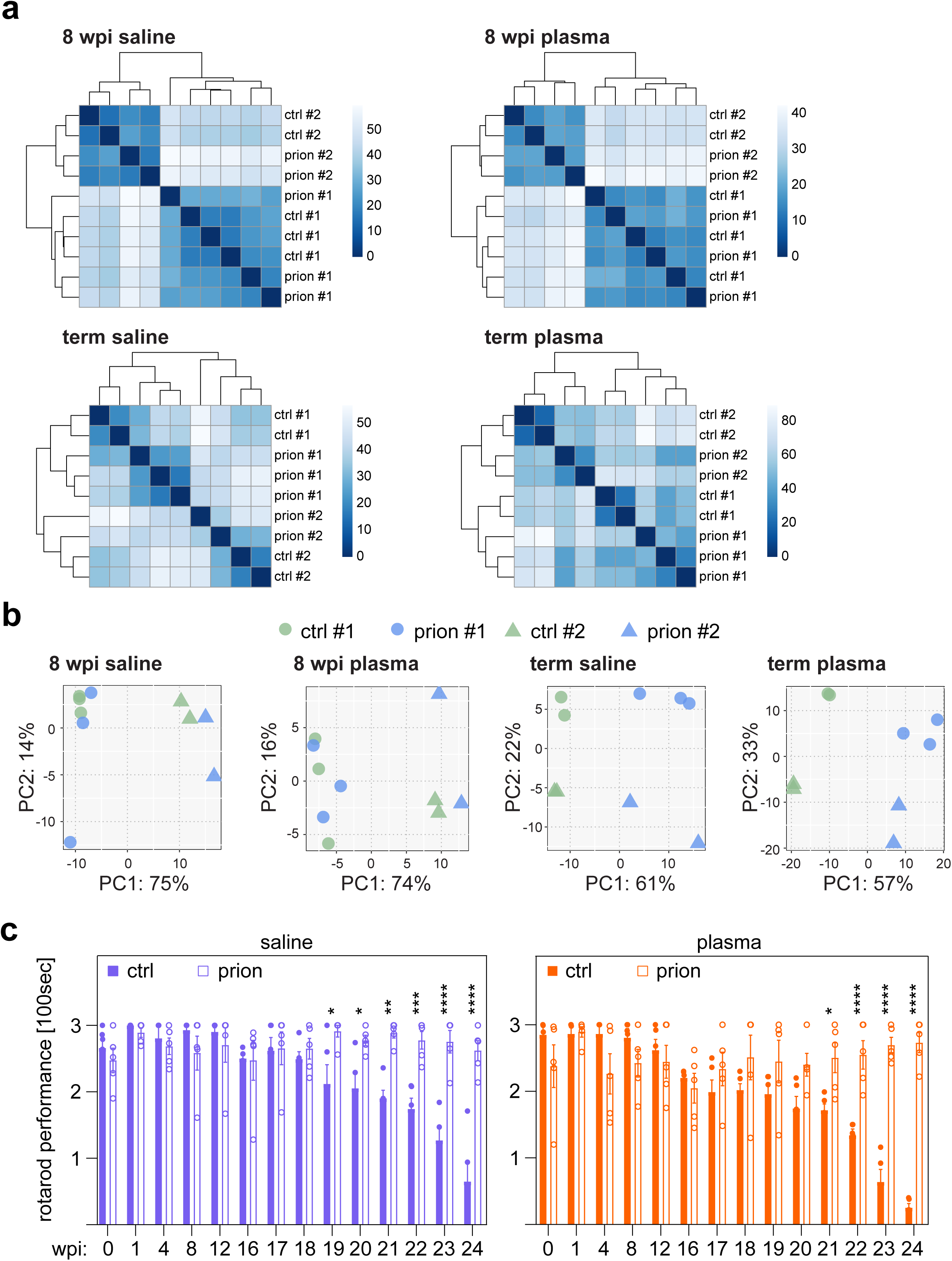
Young plasma treatment ameliorates prion disease progression. **a**, Hierarchical clustering based on Euclidean distances. Heatmaps depicting the sample distances based on RNAseq expression data. Samples cluster predominantly according to cohorts. **b**, Principal component analysis of RNAseq samples revealing a separation of control (green) and prion-injected (blue) samples at the terminal stage. Samples additionally separated according to the run of RNA isolation/processing/sequencing (run #1 versus run #2). The batch effect of the different run was accounted for during the analysis. **c**, Rotarod performance of saline (left panel) and plasma-treated (right panel) prion-inoculated mice at specified time points during disease progression (pre = pre-inoculation). Bar plots display the mean latency +/-SEM to fall in 100 seconds, with each dot representing one individual mouse. P values were calculated with a one-way ANOVA followed by Tukey’s multiple comparison test (*p<0.05; **p<0.01; ***p<0.001; ****p<0.0001; compared to 1 wpi).

## Supplementary File Legend

**Supplementary File 1: Summary table of 114 sequenced mice**

Table including sequencing and sample ID, dataset, treatment, gender, and timepoints of injection and sacrifice.

**Supplementary File 2: Expressed genes and all comparisons**

Table containing 16,560 genes that are expressed in at least one of the analyzed comparisons. Shown are GeneID, cell type enrichment if applicable (_NE = not enriched; AS = astrocytes; EC = endothelial cells; MG = microglia; N = neurons; OL = oligodendrocytes), Cluster (corresponding to clusters shown in Figure 1) and information on all edgeR comparisons of this manuscript. For each comparison expression, log_2_FC, P value and FDR are included.

**Supplementary File 3: Prion-induced gene expression changes**

Table containing 3,723 genes that are changing significantly (|log_2_FC| > 0.5 and FDR < 0.05) in at least one of the eight timepoints of the main dataset (Figure 1). Shown are GeneID, cell type enrichment if applicable (_NE = not enriched; AS = astrocytes; EC = endothelial cells; MG = microglia; N = neurons; OL = oligodendrocytes), cluster number (corresponding to clusters shown in Figure 1) and all eight edgeR comparisons related to the main dataset. For each comparison expression, log_2_FC, P value and FDR are included.

**Supplementary File 4: GO analysis of mDEGs upregulated 16wpi**

Gene Ontology analysis of terms related to biological processes, molecular function and cellular localization. 87 genes (85 of which were in the database) were upregulated from 16wpi onwards and were assessed for GO term enrichment compared to 16,127 expressed genes (15,493 of which were in the database).

**Supplementary File 5: GO analysis of mDEGs upregulated 18 wpi**

Gene Ontology analysis of terms related to biological processes, molecular function and cellular localization. 440 genes (435 of which were in the database) were upregulated from 18 wpi onwards and were assessed for GO term enrichment compared to 16,127 expressed genes (15,493 of which were in the database).

**Supplementary File 6: GO analysis of terminally downregulated DEGs**

Gene Ontology analysis of terms related to biological processes, molecular function and cellular localization. 1,081 genes (1,061 of which were in the database) were downregulated at the terminal stage and were assessed for GO term enrichment compared to 16,127 expressed genes (15,493 of which were in the database).

**Supplementary File 7: GO analysis of DEGs downregulated at 8 wpi**

Gene Ontology analysis of terms related to biological processes, molecular function and cellular localization. 813 genes (800 of which were in the database) were downregulated at 8 wpi and were assessed for GO term enrichment compared to 16,127 expressed genes (15,493 of which were in the database).

**Supplementary File 8: Differentially expressed splice isoforms**

Table containing 462 splice isoforms that are differentially expressed (FDR < 0.05) in at least one of the eight timepoints of the main dataset (Figure 4). Shown are merged, group and feature ID, genomic coordinates (chromosome, start, end, width, strand), the corresponding transcript, variant type, cell type enrichment if applicable (_NE = not enriched; AS = astrocytes; EC = endothelial cells; MG = microglia; N = neurons; OL = oligodendrocytes), cluster number (corresponding to clusters shown in Figure 4), and information on all eight DEXSeq comparisons on the main dataset. For each comparison isoform expression of ctrl and prion samples, log_2_FC, P value and FDR are included, and which isoforms are shown in Supplementary Fig. 7 is indicated. 102 splice isoforms map to genes that are differentially expressed in the main dataset. The respective RNA expression information is included for these 102 entries: cluster number (corresponding to clusters shown in Figure 1), and all eight edgeR comparisons related to the main dataset (expression, log_2_FC, P value and FDR).

**Supplementary File 9: Differentially edited sites**

Table containing differentially edited sites (p.adj < 0.05) in at least one of the eight timepoints of the main/validation dataset (Supplementary Figure 7). Shown are genomic coordinates (chromosome, start, end, width, strand), reference and alternative genomic sequence, the corresponding gene, the fraction of edited reads (of total reads) per sample and P value and adjusted P value of all eight comparisons (main and validation datasets were analyzed together).

**Supplementary File 10: Aging-related gene expression changes**

Table containing 5,613 genes that are changing significantly (|log_2_FC| > 0.5 and FDR < 0.05) in at least one of the four comparisons related to the aging dataset (8wpi_main, 8wpi_old, term_young, term_old). Shown are GeneID, cell type enrichment if applicable (_NE = not enriched; AS = astrocytes; EC = endothelial cells; MG = microglia; N = neurons; OL = oligodendrocytes), cluster number (corresponding to clusters shown in Figure 1), and information on the four edgeR comparisons related to the aging dataset. For each comparison expression, log_2_FC, P value and FDR are included.

**Supplementary File 11: Plasma-induced gene expression changes**

Table containing 3,956 genes that are changing significantly (|log_2_FC| > 0.5 and FDR < 0.05) in at least one of the four plasma dataset comparisons (8wpi_saline, 8wpi_plasma, term_saline, term_plasma). Shown are GeneID, cell type enrichment if applicable (_NE = not enriched; AS = astrocytes; EC = endothelial cells; MG = microglia; N = neurons; OL = oligodendrocytes), cluster number (corresponding to clusters shown in Figure 1), and information on the four edgeR comparisons related to the plasma dataset. For each comparison expression, log_2_FC, P value and FDR are included.

## Notes

https://histodb12.usz.ch/iMice/public/PrionRNASeqDatabase/index.php

## References

1. Aguzzi, A. & Calella, A. M. Prions: protein aggregation and infectious diseases. Physiol. Rev. 89, 1105– 1152 (2009).

2. Swerdlow, A. J., Higgins, C. D., Adlard, P., Jones, M. E. & Preece, M. A. Creutzfeldt-Jakob disease in United Kingdom patients treated with human pituitary growth hormone. Neurology 61, 783–791 (2003).

3. Collinge, J. et al. Kuru in the 21st century--an acquired human prion disease with very long incubation periods. Lancet Lond. Engl. 367, 2068–2074 (2006).

4. Prusiner, S. B. Biology and genetics of prions causing neurodegeneration. Annu. Rev. Genet. 47, 601– 623 (2013).

5. Aguzzi, A., Nuvolone, M. & Zhu, C. The immunobiology of prion diseases. Nat. Rev. Immunol. 13, 888– 902 (2013).

6. Collinge, J. Mammalian prions and their wider relevance in neurodegenerative diseases. Nature 539, 217–226 (2016).

7. Sandberg, M. K., Al-Doujaily, H., Sharps, B., Clarke, A. R. & Collinge, J. Prion propagation and toxicity in vivo occur in two distinct mechanistic phases. Nature 470, 540–542 (2011).

8. Sandberg, M. K. et al. Prion neuropathology follows the accumulation of alternate prion protein isoforms after infective titre has peaked. Nat. Commun. 5, 4347 (2014).

9. Hwang, D. et al. A systems approach to prion disease. Mol. Syst. Biol. 5, 252 (2009).

10. Majer, A. et al. Early mechanisms of pathobiology are revealed by transcriptional temporal dynamics in hippocampal CA1 neurons of prion infected mice. PLoS Pathog. 8, e1003002 (2012).

11. Xiang, W. et al. Identification of differentially expressed genes in scrapie-infected mouse brains by using global gene expression technology. J. Virol. 78, 11051–11060 (2004).

12. Majer, A. et al. The cell type resolved mouse transcriptome in neuron-enriched brain tissues from the hippocampus and cerebellum during prion disease. Sci. Rep. 9, 1099 (2019).

13. Kanata, E. et al. RNA editing alterations define manifestation of prion diseases. Proc. Natl. Acad. Sci. U. S. A. 116, 19727–19735 (2019).

14. Aguzzi, A., Lakkaraju, A. K. K. & Frontzek, K. Toward Therapy of Human Prion Diseases. Annu. Rev. Pharmacol. Toxicol. 58, 331–351 (2018).

15. Zhang, Y. et al. An RNA-Sequencing Transcriptome and Splicing Database of Glia, Neurons, and Vascular Cells of the Cerebral Cortex. J. Neurosci. 34, 11929–11947 (2014).

16. Anthony, K. & Gallo, J.-M. Aberrant RNA processing events in neurological disorders. Brain Res. 1338, 67–77 (2010).

17. Harold, D. et al. Genome-wide association study identifies variants at CLU and PICALM associated with Alzheimer’s disease. Nat. Genet. 41, 1088–1093 (2009).

18. Klebig, M. L. et al. Mutations in the clathrin-assembly gene Picalm are responsible for the hematopoietic and iron metabolism abnormalities in fit1 mice. Proc. Natl. Acad. Sci. U. S. A. 100, 8360–8365 (2003).

19. Yao, P. J., Petralia, R. S., Bushlin, I., Wang, Y. & Furukawa, K. Synaptic distribution of the endocytic accessory proteins AP180 and CALM. J. Comp. Neurol. 481, 58–69 (2005).

20. Yamada, T., Goto, I. & Sakaki, Y. Neuron-specific splicing of the Alzheimer amyloid precursor protein gene in a mini-gene system. Biochem. Biophys. Res. Commun. 195, 442–448 (1993).

21. Keren-Shaul, H. et al. A Unique Microglia Type Associated with Restricting Development of Alzheimer’s Disease. Cell 169, 1276–1290.e17 (2017).

22. Singh, M. Dysregulated A to I RNA editing and non-coding RNAs in neurodegeneration. Front. Genet. 3, 326 (2012).

23. Aguzzi, A. Prion diseases of humans and farm animals: epidemiology, genetics, and pathogenesis. J. Neurochem. 97, 1726–1739 (2006).

24. Dutta, S. & Sengupta, P. Men and mice: Relating their ages. Life Sci. 152, 244–248 (2016).

25. Villeda, S. A. et al. Young blood reverses age-related impairments in cognitive function and synaptic plasticity in mice. Nat. Med. 20, 659–663 (2014).

26. Middeldorp, J. et al. Preclinical Assessment of Young Blood Plasma for Alzheimer Disease. JAMA Neurol. 73, 1325–1333 (2016).

27. Keren-Shaul, H. et al. A Unique Microglia Type Associated with Restricting Development of Alzheimer’s Disease. Cell 169, 1276–1290.e17 (2017).

28. Iaccarino, L. et al. An In Vivo 11C-(R)-PK11195 PET and In Vitro Pathology Study of Microglia Activation in Creutzfeldt-Jakob Disease. Mol. Neurobiol. 55, 2856–2868 (2018).

29. Sorce, S. et al. The role of the NADPH oxidase NOX2 in prion pathogenesis. PLoS Pathog. 10, e1004531 (2014).

30. Pfammatter, M. et al. Absolute Quantification of Amyloid Propagons by Digital Microfluidics. Anal. Chem. 89, 12306–12313 (2017).

31. Nuvolone, M. et al. SIRPα polymorphisms, but not the prion protein, control phagocytosis of apoptotic cells. J. Exp. Med. 210, 2539–2552 (2013).

32. Dobin, A. et al. STAR: ultrafast universal RNA-seq aligner. Bioinforma. Oxf. Engl. 29, 15–21 (2013).

33. Robinson, M. D., McCarthy, D. J. & Smyth, G. K. edgeR: a Bioconductor package for differential expression analysis of digital gene expression data. Bioinforma. Oxf. Engl. 26, 139–140 (2010).

34. Goldstein, L. D. et al. Prediction and Quantification of Splice Events from RNA-Seq Data. PloS One 11, e0156132 (2016).

35. Anders, S., Reyes, A. & Huber, W. Detecting differential usage of exons from RNA-seq data. Genome Res. 22, 2008–2017 (2012).

36. Srinivasan, K. et al. Untangling the brain’s neuroinflammatory and neurodegenerative transcriptional responses. Nat. Commun. 7, 11295 (2016).

37. Nuvolone, M. et al. Strictly co-isogenic C57BL/6J-Prnp-/-mice: A rigorous resource for prion science. J. Exp. Med. 213, 313–327 (2016).

38. DePristo, M. A. et al. A framework for variation discovery and genotyping using next-generation DNA sequencing data. Nat. Genet. 43, 491–498 (2011).

39. Stilling, R. M. et al. De-regulation of gene expression and alternative splicing affects distinct cellular pathways in the aging hippocampus. Front. Cell. Neurosci. 8, 373 (2014).

